# Learning Gene Networks Underlying Clinical Phenotypes Using SNP Perturbations

**DOI:** 10.1101/412817

**Authors:** Calvin McCarter, Judie Howrylak, Seyoung Kim

## Abstract

Recent technologies are generating an abundance of genome sequence data and molecular and clinical phenotype data, providing an opportunity to understand the genetic architecture and molecular mechanisms underlying diseases. Previous approaches have largely focused on the co-localization of single-nucleotide polymorphisms (SNPs) associated with clinical and expression traits, each identified from genome-wide association studies and expression quantitative trait locus (eQTL) mapping, and thus have provided only limited capabilities for uncovering the molecular mechanisms behind the SNPs influencing clinical phenotypes. Here we aim to extract rich information on the functional role of trait-perturbing SNPs that goes far beyond this simple co-localization. We introduce a computational framework called Perturb-Net for learning the gene network that modulates the influence of SNPs on phenotypes, using SNPs as naturally occurring perturbation of a biological system. Perturb-Net uses a probabilistic graphical model to directly model both the cascade of perturbation from SNPs to the gene network to the phenotype network and the network at each layer of molecular and clinical phenotypes. Perturb-Net learns the entire model by solving a single optimization problem with an extremely fast algorithm that can analyze human genome-wide data within a few hours. In our analysis of asthma data, for a locus that was previously implicated in asthma susceptibility but for which little is known about the molecular mechanism underlying the association, Perturb-Net revealed the gene network modules that mediate the influence of the SNP on asthma phenotypes. Many genes in this network module were well supported in the literature as asthma-related.

## Introduction

One of the key questions in biology is how genetic variation perturbs gene regulatory systems to influence disease susceptibility or other phenotypes in a population. Recent advances in technologies have allowed researchers to obtain genome sequence data along with phenotype data at different levels of biological systems, such as gene expression,^1^ proteome,^2^ metabolome,^3^ and various clinical phenotype data. Combining genome sequence data with various types of molecular and clinical phenotype data in a computational analysis has the potential to reveal the complex molecular mechanisms controlled by different genetic loci that underlie diseases and other phenotypes.

To study gene regulatory systems, many previous works have considered the naturally-occurring perturbation of gene expression by genetic variants such as single nucleotide polymorphisms (SNPs), captured in expression and genotype data collected from a population. Compared to experimental perturbation methods such as gene knockdown^4^ and genome editing techniques,^5^ SNP perturbation for functional genomics studies has an advantage of being more cost effective* being easily applicable to humans, and being potentially more meaningful subtle perturbations because they exist in nature.^6^ However, it comes with the computational challenge of having to isolate the perturbation effect of each individual genetic variant, when a large number of genetic variants are perturbing the gene network simultaneously. Several computational methods have been proposed to address this challenge. Sparse conditional Gaussian graphical models (sCGGMs) have been introduced for simultaneously identifying the gene network and expression quantitative trait loci (eQTLs) from population SNP and expression data.^7,8,9^ Many other works have relied on statistically less powerful approaches of identifying eQTLs first and then incorporating the eQTLs in the network learning procedure,^10,11,12^

However, there have been relatively few works on modeling how a gene network perturbed by SNPs mediates the SNP perturbation of phenotypes. Most of the existing methods did not directly address this problem and thus, provided only limited capabilities for uncovering the molecular mechanisms behind the SNP perturbation of clinical phenotypes. Many of the previous approaches were concerned with identifying simply the co-localization of eQTLs and trait-associated SNPs,^13,14,15^ each of which were identified in a separate eQTL mapping^1,10,16,17^ and a genome-wide association study.^18,19^ These methods did not provide a description of the regulatory roles of the trait-associated SNPs beyond their co-localization with eQTLs. The genome-transcriptome-phenome structured association method^20^ focused only on identifying eQTLs and trait-associated SNPs, and was concerned with neither learning a gene network nor uncovering its role in modulating SNP effects on phenotypes. A predictive network model for diseases that involves Bayesian networks for gene regulatory networks have been proposed,^21^ but this approach relied on an elaborate pipeline of analysis to identify disease-related gene modules and genetic variants that could potentially lead to loss of statistical power.

Here, our goal is to extract rich information on the functional role of trait-perturbing SNPs that goes far beyond the simple co-localization with eQTLs, which was the focus of many of the previous studies.^13,14,15^ Towards this goal, we introduce a computational framework called Perturb-Net for directly modeling and learning the gene network that modulates the influence of SNPs on phenotypes, using SNPs as naturally occurring perturbation of a biological system. Perturb-Net builds on the key idea in the previous work on sCGGMs^7^’^8^ for learning a gene network using SNP perturbations, and models the cascade of a gene network and a phenotype netowrk under SNP perturbations as a cascade of sCGGMs, called a sparse Gaussian chain graph model (Figure 1A). Our probabilistic graphical model framework naturally leads to a set of inference algorithms for inferring a detailed description of how different parts of the gene network mediate the influence of SNPs on phenotypes, given the model estimated from population genotype, expression, and phenotype data (Figure 1B). The Perturb-Net model and inference procedures together provide a powerful tool for studying the gene regulatory mechanisms whose perturbations by SNPs lead to diseases.

**Figure 1.**
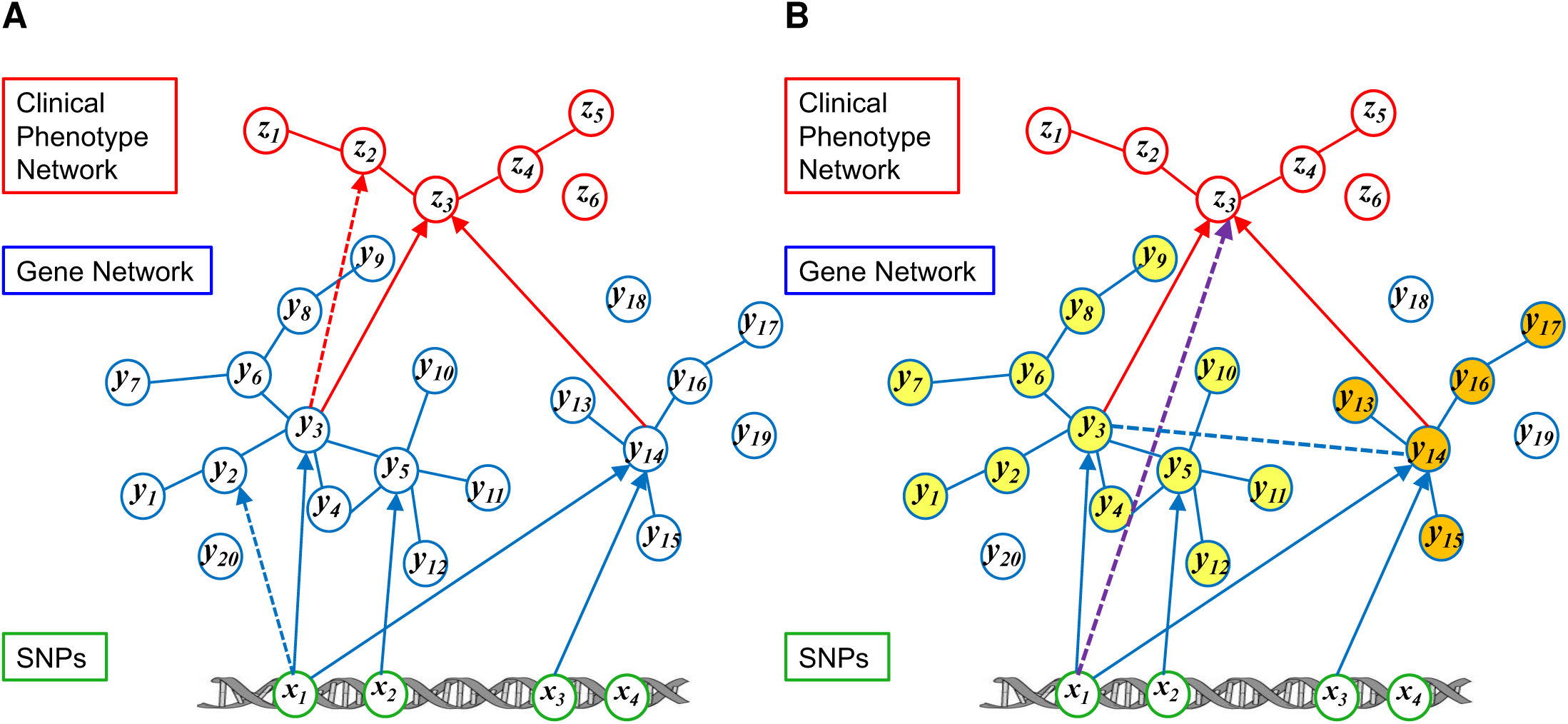
Illustration of the Perturb-Net approach. (A) Perturb-Net uses a sparse Gaussian chain graph model with a cascade of two sCGGMs, one for a gene network influenced by SNPs (blue solid edges and nodes) and the other for a clinical trait network influenced by gene expression levels (red solid edges and nodes). The sCGGM inference procedures can be used to infer hidden interactions in each of the two component sCGGMs, such as the indirect effect of SNP on expression level *y*_2_ through expression level *y*_3_ (blue dashed arrow) and the indirect effect of expression level y_3_ on phenotype through phenotype z_3_ (red dashed arrow). (B) The inference procedures of sparse Gaussian chain graph models are used to infer the information on how the gene network mediates SNP effects on phenotypes. Examples of such inferred interactions are shown for the perturbation effect of SNP *x*_1_ on phenotype *z*_3_ (purple dashed arrow), which can be decomposed into two components mediated by each of the two gene modules (yellow and orange nodes), and the inferred dependencies between expression level *y*_3_ and expression level *y*_14_ (blue dashed line) induced by phenotype *z*_3_ in the posterior gene network, after seeing the clinical phenotypes.

We present a statistically powerful and extremely efficient algorithm for learning the Perturb-Net model. The Perturb-Net learning algorithm is statistically powerful, since it estimates the entire model by solving a single optimization problem with minimal loss of statistical power and with a guarantee in finding the optimal solution due to the convexity of the optimization problem. The Perturb-Net learning algorithm is also computationally efficient and can analyze human genome-wide data with 500,000 SNPs, 11,000 gene expression levels, and several dozens of phenotype data within a few hours. The performance of the Perturb-Net learning algorithm directly depends on that of sCGGM optimization, since it uses the sCGGM learning algorithm as a key module. The previous state-of-the-art method^22^ had limited scalability due to expensive computation time and large memory requirement, requiring more than 4 hours for only 10,000 SNPs and running out of memory for 40,000 SNPs. We present a new learning algorithm Fast-sCGGM and its extension Mega-sCGGM with orders-of-magnitude speed-up in computation time that runs on a single machine without running out of memory and that is parallelizable. Our new sCGGM learning algorithms allow Perturb-Net to be applied to human genome-wide data.

We demonstrate Perturb-Net on the data collected for participants in the Childhood Asthma Management Program (CAMP).^23,24,25^ Perturb-Net revealed the asthma gene network and how different parts of this gene network mediate the SNP perturbations of phenotypes. Furthermore, for a locus that was previously implicated in asthma susceptibility but for which little has been known about the molecular mechanism underlying the association, Perturb-Net revealed the gene network modules that mediate the influence of the SNP on asthma phenotypes. Many genes in this network module were well supported in the literature as asthma-related, suggesting our framework can reveal the molecular mechanisms underlying the SNP perturbations of phenotypes.

## Material and Methods

learning/inference algorithms for Perturb-Net for learning the gene network under SNP perturbations that underlies clinical phenotypes.

### Perturb-Net model

Let x ∈ {0,1,2}^*p*^ denote minor allele frequencies at *p* loci of an individual, y ∈ ℝ^*q*^ expression levels for *q* genes, and z ∈ ℝ^*r*^ measurements for *r* phenotypes. Then, Perturb-Net models the cascaded influence of SNPs on a gene network and a phenotype network as a Gaussian chain graph model (Figure 1A), which is a factorized conditional probability distribution defined as follows:

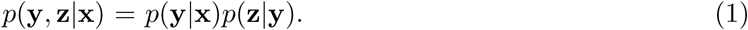

Each probability factor above is modeled as a conditional Gaussian graphical model (CGGM): ^7,8,22^

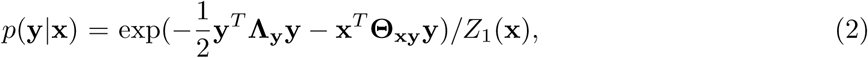

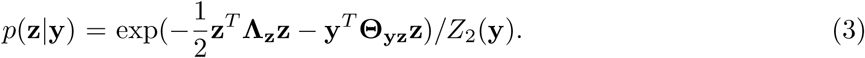

The first probability factor in Eq. (2) models the gene network perturbed by SNPs, representing the gene network as a *q × q* positive definite matrix Λ_y_ and the SNP perturbation of this network as Θ_xy_ ∈ ℝ^*p*×*q*^. The second probability factor in Eq. (3) models the phenotype network Λ_z_, a *r*×r positive definite matrix, and the perturbation of this network by gene expression levels Θ_xy_ ∈ ℝ^*q*×*r*^. The *Z*_1_(x) and *Z*_2_(y) in Eqs. (2) and (3) are the constants for ensuring that each CGGM is a proper probability distribution that integrates to one. Our model in Eq. (1) defines a probability distribution over the graph shown in Figure 1A. Thus, a non-zero value in the *(i, j*)th element of the network parameters, |Λ_y_]_*i,j*_ of Λ_y_ and [Λ_z_]_*i,j*_ of Λ_z_, corresponds to presence of an edge between the *i*th and *j*th expression or clinical phenotypes. Similarly, a non-zero value in the (*i,j*)th element of the perturbation parameters, [Θ_xy_]_*i,j*_ of Θ_xy_ and [Θ_yz_]_*i,j*_ of Θ_yz_, indicates an edge between the *i*th perturbant and the *j*th expression or clinical phenotype.

This Gaussian chain graph model corresponds to the continuous counterpart of the chain graph model obtained by threading conditional random fields (CRFs) for discrete random variables. CRFs and the chain graph models built from CRFs have been hugely popular in other application areas of statistical machine learning such as text modeling and image analysis for modeling multiple correlated output features influenced by input features. ^26,27,28^ Here, we explore the use of a chain graph model constructed with sCGGMs, corresponding to Gaussian CRFs, and develop an extremely fast learning algorithm that runs on human data within a few hours and a set of inference algorithms for dissecting the gene regulatory mechanisms that govern the influence of SNPs on phenotypes.

### Perturb-Net learning algorithms

We present an extremely efficient algorithm for obtaining a, sparse estimate of the model parameters with few edges in the graph. The Perturb-Net learning algorithm minimizes the negative log-likelihood of data with *L*_1_ regularization,^29^ which is a convex optimization problem with a guarantee in finding the optimal solution. Given genotype data **X** ∈ ℝ^*n*×*p*^ for *n* samples and *p* SNPs, expression data **Y** ∈ ℝ^*n*×*q*^ for *q* genes, and phenotype data **Z**∈ ℝ^*n*×*r*^ for *r* phenotypes for the same *n* samples, we estimate a sparse Gaussian chain graph model in Eq. (1) by minimizing the negative log-likelihood of data along with sparsity-inducing *L*_1_ penalty:

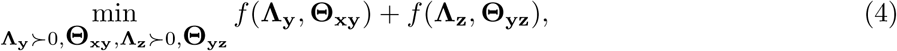

Where

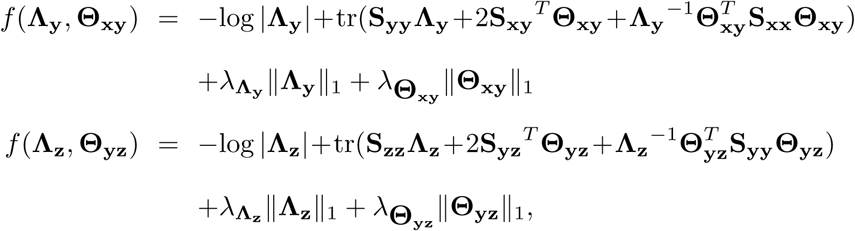

given data covariance matrices 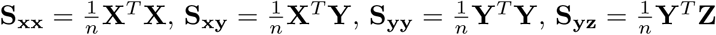,and 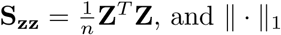, and || · ||_1_ for the non-smooth elementwise *L*_1_ penalty. The regularization parameters 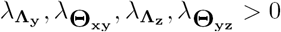are chosen to maximize the Bayesian information criterion (BIC). We do not penalize the diagonal entries of **Λ**_y_ and **Λ**_z_, following the common practice for sparse inverse covariance estimation.

The above optimization problem decouples into two subproblems, each containing one of two disjoint sets of parameters {**Λ**_y_, **Θ**_xy_} and {**Λ**_z_, **Θ**_yz_}, each of which can be solved with an sCGGM optimization algorithm. Since our learning algorithm uses an sCGGM learning method as a key module, we developed sCGGM learning algorithms called Fast-sCGGM for reducing computation time and Mega.-sCGGM for futher improving Fast-sCGGM to remove the memory constraint, both of which are parallelizable over multiple cores of a machine. Fast-sCGGM improves the computation time of the previous method by alternately optimizing the network parameter (**Λ**_y_ and **Λ**_z_) and the perturbation parameters (**Θ**_xy_ and **Θ**_yz_), where each of the alternate optimization can be efficiently solved using the fast Lasso optimization technque as the key subroutine (see Appendix A for detail).

While Fast-sCGGM improves the computation time of the previous method, it is limited by the memory size required to store large *q*×*q* or *p*×*q* matrices during the iterative optimization. A naive approach to reduce the memory footprint would be to recompute portions of these matrices on demand for each coordinate update, which would be very expensive. Hence, we combine Fast-sCGGM with block coordinate descent and introduce the Mega-sCGGM algorithm to scale up the optimization to very large problems on a machine with limited memory. During the iterative optimization, we update blocks of the large matrices so that within each block, the computation of the large matrices can be cached and re-used. These blocks are determined automatically in each iteration by exploiting the sparse stucture (see Appendix A for detail).

We introduce a modification of our learning algorithm for semi-supervised learning, to handle the situation where expression data are available only for a subset of individuals because of the difficulty of obtaining tissue samples. This modification corresponds to an expectation maximization (EM) algorithm^30^ that imputes the missing expression levels in the E-step and performs our Fast-sCGGM or Mega-sCGGM optimization in the M-step. For semi-supervised learning, given a dataset 𝒟= {𝒟_o_, 𝒟_h_}, where𝒟 _o_ = {X_o_,Y_o_,Z_o_} for the fully-observed data and 𝒟*h* = {X*h*,Z*h*} for the samples with missing gene-expression levels, we adopt an EM algorithm^30^ that iteratively maximizes the expected log-likelihood of data:

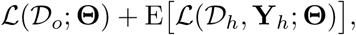

combined with *L*_1_-regulariza,tion, where *ℒ*(𝒟_0_;**Θ**) and *ℒ*{𝒟*f*_h_;**Θ**) are the log-likelihood of data 𝒟_0_ and 𝒟_h_ with respect to the model in Eq. (1) and the expectation is taken with respect to:

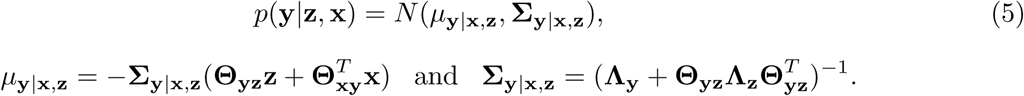

A naive implementation of this EM algorithm leads to an algorithm that requires expensive computation time and large storage of dense matrices that exceeds the computer memory. To make the EM algorithm efficient in terms of both time and memory, we embed the expensive E-step computation within the M-step, using a low-rank representation of dense matrices (see Appendix B). This implementation produces the same estimate as the original EM algorithm.

### Perturb-Net inference procedures

While the sparse Gaussian chain graph model explicitly represents pair-wise dependencies among variables as edges in the graph, there are other dependencies that are only implicitly represented in the model but can be revealed by performing an inference on the estimated probabilistic graphical model. Here, we provide an overview of the inferred dependencies, all of which involve simple matrix operations.

The following two inference methods directly follow from the inference method for an sCGGM (Figure 1A),^7,8^ which infers the indirect perturbation effects that arise from the direct perturbation effects propagating to other parts of the network.

- **Indirect SNP perturbation effects on gene expression levels: B_xy_ =−Θ_xy_Λ_y_^-1^** where [**B**_xy_|*j:sj*_*i,j*_ represents the indirect perturbation effect of SNP *i* on the expression level of gene *j* (blue dashed arrow in Figure 1A). This can be seen by deriving the marginal distribution from the sCGGM component model *p*(**y|x**) as follows:

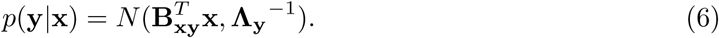 From Eq. (6), the marginal distribution for the expression level [y]_*i*_ of gene *i* can be obtained as *p*([y]_*i*_|x) = *N*([[**B**_xy_]_:,>*i*_]^*T*^ x,[Λ_y_^-1^]_*i,j*_) While [**Θ** [**0_xy_**]_*i,j*_. represents the direct perturbation effect of SNP *i* on the expression of gene *j*, [B_xy_];. _*i,j*_. represents the overall perturbation effect that aggregates all indirect influence of this SNP on gene *j* through other genes. Wien SNP *i* does not influence the expression of gene *j* directly but exerts influence on gene *j* through other genes connected to gene *j* in the network **Λ**_y_, we have. [**Θ_xy_**]_*i,j*_)=0 but [B**_xy_**]_*i,j*_≠0.
- **Indirect effects of gene expression levels on clinical phenotypes: B_yz_ = 0-Θ_yz_Λ ^-1^**. where [**B**_yz_]_*i,j*_ represents the indirect influence of the expression level of gene *i* on phenotype *j* (red dashed arrow in Figure 1A). Similarly as above, this can be seen by deriving the marginal distribution from the sCGGM component model *p*(z|y), as follows:

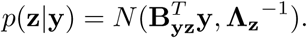 Then, the marginal distribution for [**z**]_*i*_ of phenotype *i* can be obtained as *p*([**z**]_*i*_|**y**) =*N*([[B_yz_]:,*i*]^*T*^y, [Λ_z_^-1^]_*i,j*_). While [**Θ_yz_**]_*i,j*_] represents the direct influence of gene *i* on phenotype *j*, [B_yz_] _*i,j*_. represents the overall influence that aggregates all indirect influence of this expression level on phenotype *j* through other phenotypes. The sparse Gaussian chain graph model provides the following additional inference procedures for extracting the information on whether SNP perturbation effects on the gene network reach the phenotypes and how different genes or subnetworks of the gene network mediate SjtP effects on phenotypes (Figure 1B).
- **SNP effects on clinical phenotypes:** B_xz_ = B_xy_B_yz_, where [B_xz_]_*i, j*_ represents the overall influence of SNP *i* on phenotype *j* mediated by gene expression levels in gene network Λ_y_ (purple dashed arrow in Figure 1B). The effects of SNPs on phenotypes are not directly modeled in our model but can be inferred by deriving the marginal distribution *p*(**z**|**x**) as follows:

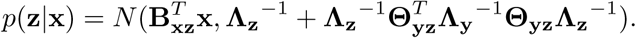 The marginal distribution for the phenotype [z]_*i*_ of phenotype *i* given x can be obtained as 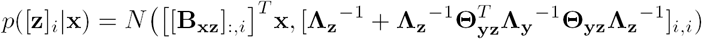, where each element [**B**_xz_]_*i,j*_ represents the overall influence of SNP *i* on phenotype *j* mediated by the gene network in **Λ_y_** and other phenotypes connected to phenotype *j* in **Λ_z_**.
- **SNP effects on clinical phenotypes mediated by a gene module:** The overall SNP effects on phenotypes in **B**_xz_ above can be decomposed into the SNP effects on phenotypes mediated by each gene module. Let *M* be a gene module that consists of a subset of the *q* genes whose expression levels were modeled in **Λ**_y_ (yellow and orange gene modules in Figure1B). Then, the effects of SNPs on phenotypes mediated by the genes in module *M* can be obtained as follows:

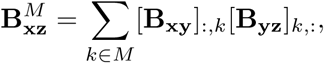

where [**B**_xy_]_:,a_ o represents the ath row of **B**_xy_ and [**B**_yz_]_b,:_ represents the 6th column of **B**_yz_. In the above equation,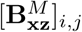 j quantifies the effect of SNP *i* on phenotype *j* through the expression levels of genes in module *M*. If *M*_1_,…, *M*_s_ are disjoint subsets of *q* genes, where ∪_m=1,.…,,S_*M*_m_ is the full set of *q* genes, we have the following decomposition:

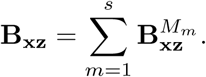
- **Inferred dependencies among genes after seeing phenotype data:** 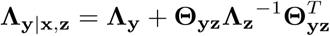 represents gene network **Λ**_y_ augmented with the component 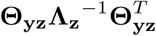 introduced through dependencies in phenotype network **Λ**_z_ (blue dashed edge in Figure 1B). In this augmented network, additional edges are introduced between two genes if their expression levels influence the same trait or if they both affect traits that are connected in the phenotype network **Λ**_z_. The posterior gene network **Λ**_y_|_x z_, which contains the dependencies among expression levels after taking into account phenotype data, can be obtained by inferring the posterior distribution given phenotypes from the estimated Gaussian chain graph model as follows:

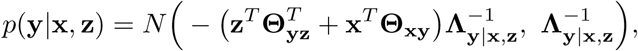

where

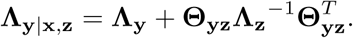 The inferred network **Λ**_y|x,z_ can also be seen by inferring from the estimated model the joint distribution

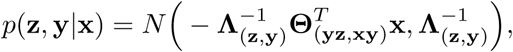

where **Θ**,_(yz, xy)_ = (0_P × r_, **Θ**_xy_) with *p* x× *r* matrix of 0’s and Λ (z y*j*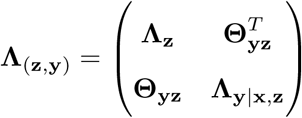. This joint distribution is an alternative representation of the same Gaussian chain graph model in Eq. (1) and corresponds to another sCGGM over **y** and **z** conditional on **x.** This process of introducing the additional dependencies via 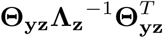 in this new sCGGM, which is equivalent to the original chain graph model, is also known as moralization in the probabilistic graphical model literature.^27^

### Prediction tasks

We use the estimated Perturb Net model and the results of probabilistic inference on this model to make predictions on previously unseen patients. From each of the two component sCGGMs in our model, we make the following predictions:

- 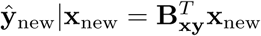for predicting the expression levels ŷ_new_ given the genotypes x_new_ of a new patient
- 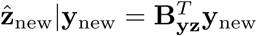for predicting the phenotypes 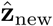 given the expression levels **y**_new_ of a new patient

From the full sparse Gaussian chain graph model, we make the following predictions:

- 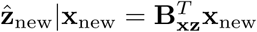 for predicting the phenotypes 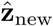 given the genotypes **x**_new_ of a new patient
- 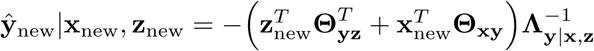for predicting the expression levels ŷ_new_ given the genotypes x_new_ and the phenotypes x_new_ of a new patient

### Lasso for comparison with our algorithms

We compare the performance of our method with that of Lasso,^29,31^ a popular statistical method based on linear regression models for studying the associations among SNPs, expression measurements, and phenotypes. We begin by setting up a two-layer multivariate regression model for genotypes x ∈ {0,1, 2} ^*p*^, expression measurements ∈ℝ^*q*^, and phenotypes z G R∈ℝ ^r^ as follows:

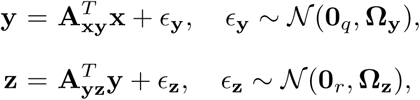

where **A**_xy_ ∈ℝ^*p* × *q*^ and **A**_yz_ ∈ℝ^*q*×*r*^ are regression coefficients, *ϵ*_y_ ∈ℝ^*q*^and *ϵ*_z_ ∈ℝ^*r*^z are noise distributed with zero means and diagonal covariances 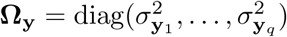 and 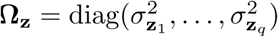.

Given genotype data **X** ∈ {0,1,2}^*n*×*p*^ for *n* samples and *p* SNPs, expression data Y ∈ℝ^*n*×*q*^ for *q* genes, and phenotype data **Z**∈ℝ^*n*×*r*^ G R"f for *r* phenotypes, we obtain a Lasso estimate of the regression coefficients by minimizing *L*_1_-regularized negative log-likelihood as follows:

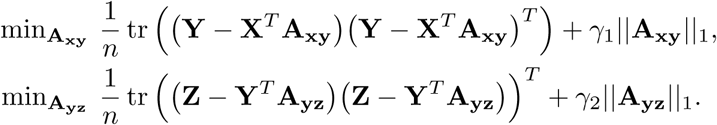

Using the Lasso estimate of the regression coefficients **A**_xy_ and **A**_yz_, we compute predictions for this model analogously to our sparse Gaussian chain graph model.

- 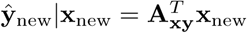
- 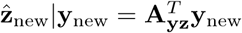
- 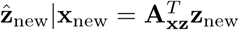
- 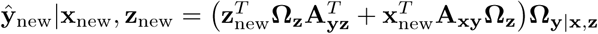 where **Ω**_y|x,z_ = ([**Ω**_y_]^-1^ +**A**_yz_[**Ω**_y_]^-1^**A**_yz_^*T*^)^-1^. For this prediction task, we estimate the variances as follows:^32^

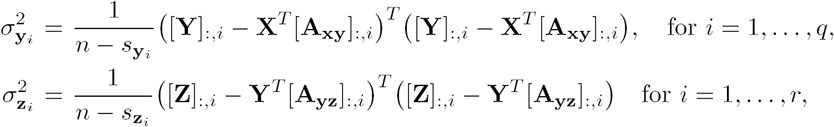

where 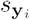 and 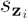 are the numbers of non-zero entries in [**A**_xy_]_;,*i*_ and [**A**_yz_]:_,*i*_ respectively.

### Preparation of asthma dataset

We applied our method to a dataset comprising genotype., gene expression, and clinical phenotype data, collected from asthma patients participating in CAMP study.^23,24,25^ We used 174 non-Hispanic Caucasian subjects for whom both genotype and clinical phenotype data were available. For a subset of 140 individuals, gene expression data from primary peripheral blood CD4+ lymphocytes were also available. After removing SNPs with minor allele frequency less than 0.1 and those with missing reference SNP ids, we obtained 495,597 SNPs for autosomal chromosomes. We then imputed missing genotypes using fastPHASE.^33^ Given expression levels for 22,184 mRNA transcripts profiled with Illumina HumanRef8 v2 BeadChip arrays,^25^ we removed transcript levels with expression variance less than 0.01, which resulted in a set of 11,598 expression levels to be used in our analysis. Then, we converted the expression values to their *z*-scores. The clinical phenotype data comprised 35 phenotypes (Table S1), including 25 features related to lung function and 10 features collected via blood testing.. The clinical phenotypes were converted to their *z*-scores within each phenotype so that all phenotypes have equal variance. We then imputed missing values using low-rank matrix completion.^34^

### Comparison of the computation time of different algorithms

In order to compare the computation of different algorithms, we used the following software and hardware setup. For Lasso, we used the implementation in GLMNET^35^ with a backend written in Fortran. For Newton coordinate descent, which is the previous state-of-the-art approach for optimizing sCGGMs, we took the implementation written in C++ provided by the authors^22^and sped up this implementation with the Eigen matrix library, by employing low-rank matrix representations and using sparse matrix multiplications. For all methods, the code was compiled and run with OpenMP multi-threading enabled on the same machines with 20Gb of memory and 16 cores. We used the same regularization parameters for our method and the previous method for sCGGM optimization, so the resulting solutions were identical with the same sparsity levels. For Lasso, we chose the regularization parameters so that the *L*_1_-norm of the regression matrix roughly matched that of our inferred indirect SNP effects.

## Results

### Comparison of the scalability of Mega-sCGGM and other methods

We assess the scalability of Mega-sCGGM and other previous algorithms on the expression measurements of 11,598 genes and the genotypes of 495,597 SNPs for 140 subjects from the CAMP data. We estimated sCGGMs, using both our new method and the previous state-of-the-art method based on the Newton coordinate descent method..^22^ Since the sCGGM optimization problem is convex with a single globally optimal solution, both our and previous methods obtain the same parameter estimates, although the computation time differs between the two methods. We also obtained the computation time of Lasso implemented in GLMNET, ^29,35^ the well-known computationally efficient algorithm for learning a simple but less powerful regression model. Although the sparse multivariate regression with covariance estimation^36^ has also provided a methodology that could be used for learning a gene network influenced by SNPs, this approach has been found to take days to learn a model from a small dataset of only 1,000 SNPs and 500 gene expression levels,^8^ so we did not include it in our experiment, All of the optimization methods were run on the same hardware setup with comparable software implementations.

In our comparison of different methods, our algorithm significantly outperformed the previous state-of-the-art method for learning an sCGGM in terms of both computation time and memory requirement and scaled similarly to Lasso (Figure 2). In comparison of our method with Lasso on datasets with 40,056 SNPs from chromosome 1, 21,757 SNPs for chromosomes 1 through 6, and 495,597 SNPs from all autosomal chromosomes and all expression measurements, our method was not substantially slower than Lasso, even though our method learns a more expressive model than Lasso. The previous sCGGM optimization algorithm ran out of memory even on the smallest dataset above with SNPs only from chromosome 1, so we compared the two algorithms on a much smaller dataset with 1,000 and 10,000 SNPs. On 10,000 SNPs, the previous algorithm for sCGGM required more than four hours, whereas in less than four hours, our algorithm was able to run on all 495,597 SNPs.

**Figure 2.**
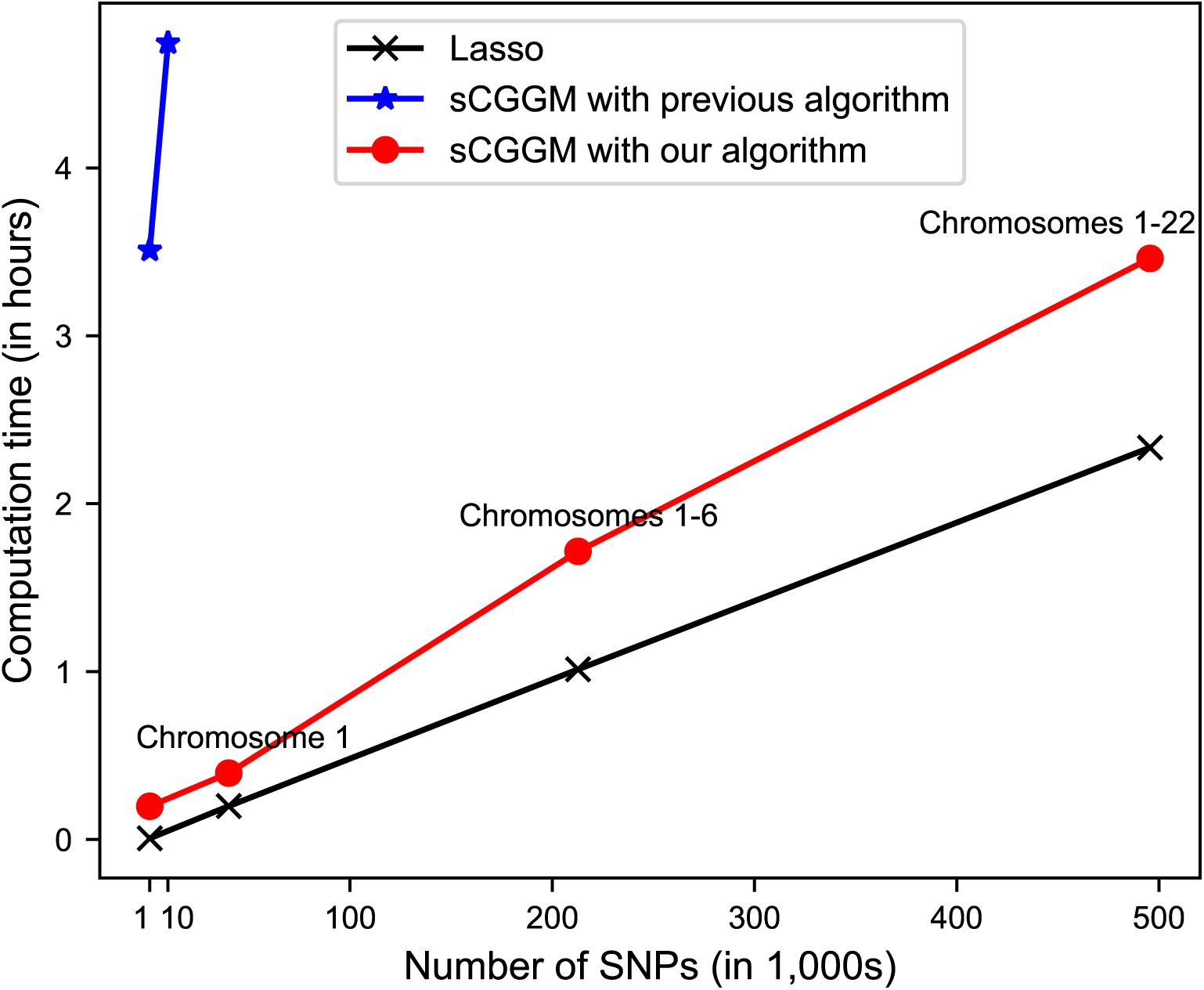
Comparison of computation time of different methods. The computation time of our Mega-sCGGM is compared with that of previous learning algorithm for sCGGMs and Lasso. We applied all methods to all expression data and genotype data from chromosome 1, chromosomes 1-6, chromosomes 1-16, and chromosomes 1-22. The previous algorithm for sCGGMs ran out of memory at chromosome 1, so we obtained its computation time with much smaller datasets with 1,000 and 10,000 SNPs.

### Analysis of asthma data

We now fit a sparse Gaussian chain graph model to the genotype, expression, clinical phenotype data gathered from participants in the Childhood Asthma Management Program (CΛMP).^23,24,25^ Λfter preprocessing the data, we applied our method to the data from f40 subjects for whom all data were available for 495,597 SNPs on 22 autosomal chromosomes, 11,598 gene expression levels, and 35 phenotypes (Table S1) and 34 additional subjects for whom data were available only for genotypes and phenotypes but not for expression levels. Below we perform a detailed analysis of the estimated model.

#### Overview of the Perturb-Net model

We first examined the overall estimated model for the module structures in the phenotype and gene networks (Figure 3). To see the structure in the phenotype network **Λ_z_**, we reordered the nodes of the network by applying hierarchical clustering to each set of the lung function and blood test phenotypes. This revealed the dense connectivities within the two known groups of phenotypes and the two sub-clusters within the group of lung function phenotypes (Figure 3A).

**Figure 3.**
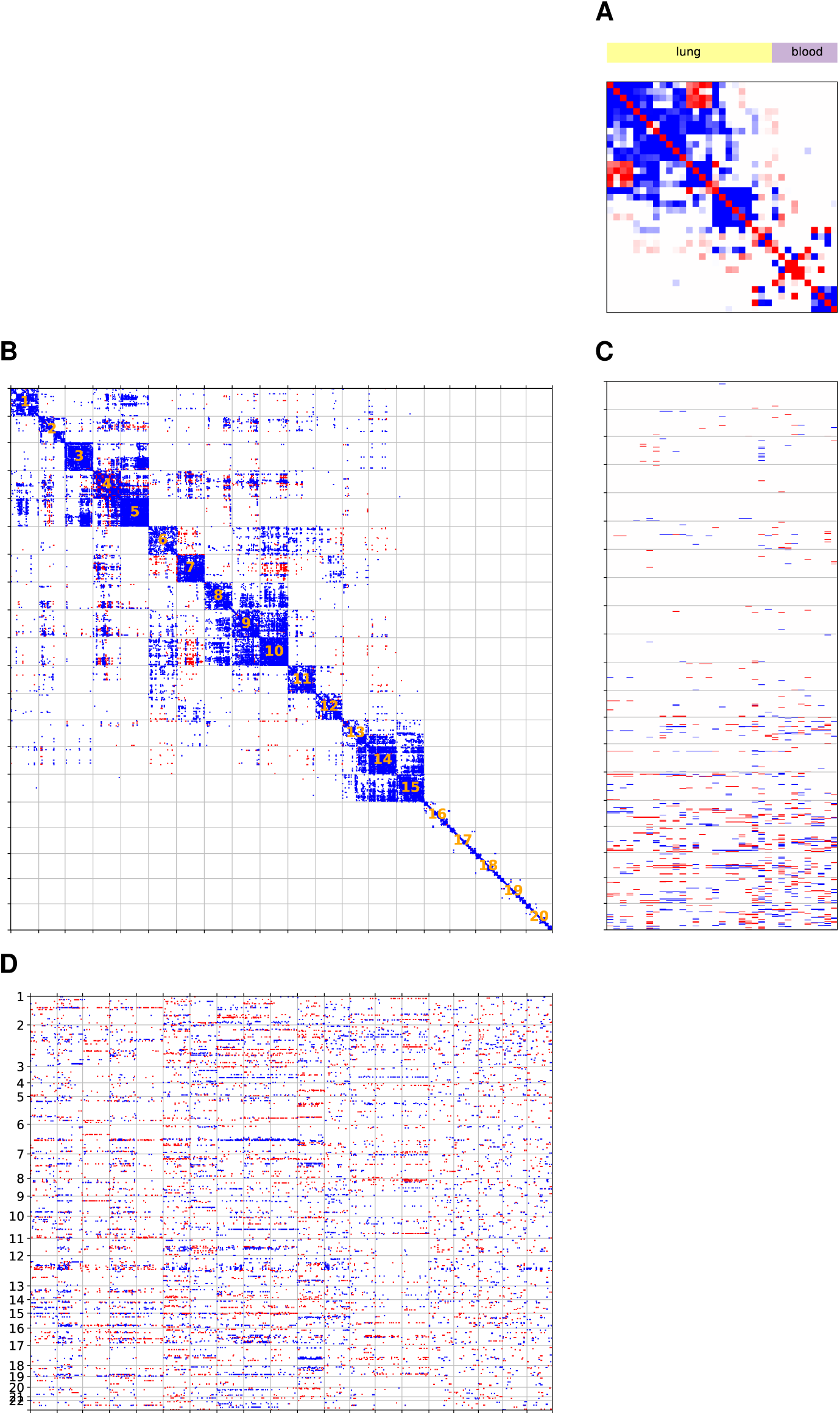
The Perturb-Net model estimated from asthma data. The parameters of the sparse Gaussian chain graph model estimated from the asthma data are shown. (A) Asthma phenotype network **Λ_z_.** The phenotypes were ordered by hierarchical clustering applied to within each of the two groups of phenotypes, lung function traits (yellow) and blood test traits (purple). (B) Gene network **Λ_y_,** The gene network is annotated with 20 modules obtained from applying a network clustering algorithm METIS^37^ to **Λ_y_.** (C) The influence of gene expression levels on phenotypes Θ_yz_. (D) SNP perturbation of gene expression levels Θ_xy_ for the top 1,000 eQTL hotspots, ordered by genomic location and labeled by chromosomes. In each panel, non-zero elements of the estimated parameters are shown as blue for positive interactions and red for negative interactions.

The gene network **Λ_y_** also showed a clear module structure (Figure 3B). To find the module structure in the network, we identified the genes tha.t are connected to at least one other gene in the network **Λ_y_**and partitioned the network over those genes into 20 subnetworks with roughly equal number of nodes, using the network clustering algorithm METIS.^37^ Out of 11,598 genes,6,102 genes were connected to at least one other gene in the network. For the rest of our analysis, we focus on the network and modules over the 6,102 genes, since these genes are likely to form modules for pathways with a functional impact on asthma phenotypes. Modules 1-15 were densely connected clusters of co-expressed genes, suggesting those modules are likely to consist of a functionally coherent set of genes, whereas modules 16-20 had relatively fewer edge connections within each cluster.

Next, we considered the effects of the gene modules on the lung and blood phenotypes in Θ_yz_ and the SNP perturbations of the gene modules in Θ_xy_. Modules 1-12 had relatively small effects on the phenotypes despite their dense connectivities, whereas modules 13-20 appeared to have stronger effects on both groups of phenotypes (Figure 3C). The SNP effects on the modules in Θ_xy_ for the top1000 eQTL hotspots, determined by overall SNP effects on all genes (Σ_*j*_|[Θ_xy_]_*i,j*_| for each SNP *i*), showed that many of these hotspots perturb the expression of genes in the same module in the gene network (Figure 3D). Given these observations from the visual inspection of Θ_yz_ and Θ_xy_, we summarized Θ_yz_ and Θ_xy_ at module level and compared the module-level summaries across modules. To quantify the module-level influence of expression levels on each group of phenotypes, from the direct influence Θ_yz_ and indirect influence B_yz_ we computed the overall effect sizes of all genes in the given gene module on all phenotypes in each of the lung and blood phenotype groups (Σ_*i*∈M,*j*∈K_|[**Θ**_yz_]_*i,j*_| and Σ_*i*∈M,*j*∈K_|[**B**_yz_]_*i,j*_| for each gene module *M* and phenotype group *K*). Similarly, from **Θ** 0_xy_ and **B**_xy_ we computed the overall SNP effect sizes on all genes in the given module (Σ_*i,j*∈M_|[**Θ**_yz_]_*i,j*_| and Σ_*i,j*∈M_|[**B**_XY_]_*i,j*_|for each SNP *i* and module *M).*

Among the 20 20 gene modules, modules 13-20 overall had stronger influence on both lung and blood phenotypes than the other gene modules (Figures 4A and 4B), although SNP perturbations were found across all gene modules without any preference to those modules with stronger influence on phenotypes (Figure 4C). For modules 13-20, the overall effect sizes on the lung phenotypes ranged between 0.8 and 7.5 for direct and indirect influence with an exception of module 14, whereas for modules 1-12, the overall effect sizes were less than 0.8 (Figure 4A). Modules 13-20 also had strong effects on the blood phenotypes (Figure 4B), although module 13 had substantially stronger effect on the blood phenotypes than on the lung phenotypes. On the other hand, the overall SNP effects were similar across all gene modules for both the direct and indirect SNP effects (Figure 4B). The overall indirect SNP effects were larger for some modules (e.g., module 14), but this was largely because of the substantially stronger edge connectivities in that module, which led to stronger propagation of the direct SNP perturbation effects.

**Figure 4.**
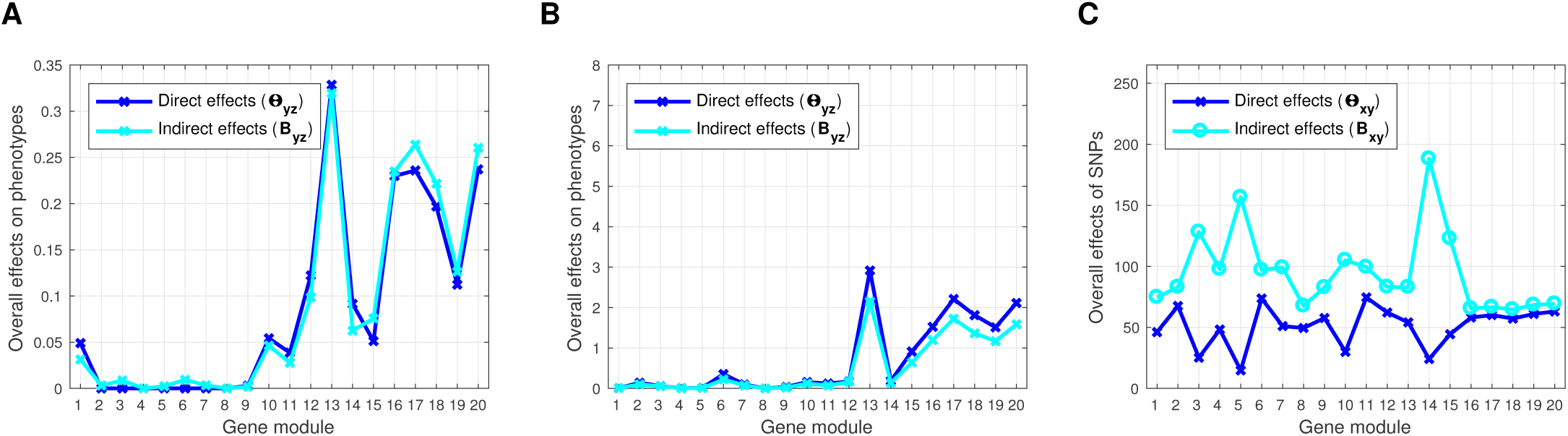
SNP effects on gene modules and gene-module effects on phenotypes. Given the estimated **Θ**_yz_ and inferred **B**_yz_ for gene-expression effects on phenotypes from the Perturb-Net model, we show the gene-module effects on each group of phenotypes for (A) lung and (B) blood, computed as the sum of absolute effect sizes across all genes within the module and across all phenotypes in the group..(C) Given the estimated **Θ**_xy_ and inferred **B**_xy_ for SNP effects on gene network, we show the SNP effects on each gene module, summarized as the sum of absolute effect sizes across all SNPs and all genes within the module.

#### Gene modules that influence phenotypes are enriched for immune genes

To determine the functional role of the gene modules, we performed gene ontology (GO) gene set enrichment analysis.^38,39^ For each module, we performed a Fisher’s exact test to find the significantly enriched GO categories in biological processes (*p*-value < 0.05 after Bonferroni correction for multiple testing), using the GO database with annotations for 21,002 genes.

Among all 20 modules, modules 13-15 had a statistically significant enrichment of GO terms related to immune system function, which also corresponded to the most significant enrichments across all modules (Table 1). Even though modules 16-20 did not have any significant enrichment of asthma-related GO categories, many of the genes in these modules were connected to genes in modules 13-15 in the posterior gene network **Λ**_y|*i*X),Z_ (Figure 5), and thus this subset of genes in modules 16-20 may be also involved with immune system function. To see if this is indeed the case, we obtained the significantly enriched GO categories in the 374 genes in modules 16-20 that are connected to modules 13-15 in the posterior network **Λ**_y|X),Z_ (*p*-value < 0.05 after Bonferroni correction). This set of genes was significantly enriched for several GO categories rela.ted to immune system processes, including cellular response to stress (*p*-va.lue = 2.90× Xl0 ^-2^ with overlap of 35 genes out of 1599 genes in the category), regulation of defense response to virus (*p*-value = 4-36 ×l0^-2^ with overlap of 6 genes out of 71 genes in the category), and regulation of immune effector process (*p*-value = 4.42 ×10^-2^ with overlap of 14 genes out of 409 genes in the category).

**Table 1.** GO categories enriched in gene modules in the estimated asthma gene network

|  | Size | Biological Process Pathway | P-value | Overlap* |
| --- | --- | --- | --- | --- |
| 1 | 314 | Cellular macromolecule metabolic process | $2.94 \times 10^{-12}$ | 159 / 7006 |
| 2 | 297 | Cellular nitrogen compound metabolic process | $1.64 \times 10^{-4}$ | 112 / 5164 |
| 3 | 314 | Nucleobase-containing compound metabolic process | $5.62 \times 10^{-13}$ | 123 / 4538 |
| 4 | 314 | Organelle organization | $2.02 \times 10^{-4}$ | 79 / 3167 |
| 5 | 314 | Nucleobase-containing compound metabolic process | $4.58 \times 10^{-4}$ | 106 / 4538 |
| 6 | 314 | Unclassified | NA | NA |
| 7 | 314 | Cellular localization | $1.80 \times 10^{-4}$ | 60 / 2287 |
| 8 | 314 | Cellular metabolic process | $1.73 \times 10^{-4}$ | 178 / 9003 |
| 9 | 314 | Cellular metabolic process | $8.39 \times 10^{-6}$ | 185 / 9003 |
| 10 | 314 | Macromolecule metabolic process | $8.73 \times 10^{-5}$ | 159 / 7749 |
| 11 | 314 | Heterocycle metabolic process | $7.63 \times 10^{-3}$ | 103 / 4715 |
| 12 | 298 | Translation | $1.65 \times 10^{-9}$ | 27 / 383 |
| 13* | 297 | Immune system process | $3.66 \times 10^{-12}$ | 83 / 2552 |
| | | Response to stimulus | $1.70 \times 10^{-9}$ | 162 / 8009 |
| | | Response to stress | $8.43 \times 10^{-9}$ | 90 / 3333 |
| 14* | 314 | Immune response | $3.96 \times 10^{-38}$ | 106 / 1673 |
| | | Leukocyte activation involved in immune response | $9.04 \times 10^{-31}$ | 62 / 607 |
| | | Granulocyte activation | $1.24 \times 10^{-26}$ | 53 / 495 |
| 15* | 313 | Cell activation in immune response | $4.95 \times 10^{-32}$ | 65 / 611 |
| | | Myeloid leukocyte activation | $5.28 \times 10^{-32}$ | 63 / 566 |
| | | Immune system process | $1.01 \times 10^{-31}$ | 124 / 2552 |
| 16 | 290 | Cellular process | $3.73 \times 10^{-3}$ | 132 / 15013 |
| 17 | 290 | Regulation of macromolecule metabolic process | $1.58 \times 10^{-3}$ | 74 / 6142 |
| 18 | 282 | Organonitrogen compound metabolic process | $1.97 \times 10^{-2}$ | 60 / 5523 |
| 19 | 285 | Cell cycle | $1.50 \times 10^{-8}$ | 37 / 1355 |
| 20 | 295 | Cellular component organization or biogenesis | $4.34 \times 10^{-3}$ | 66 / 5525 |

\* The number of genes in the overlap / the total number of genes in the GO category

**Figure 5.**
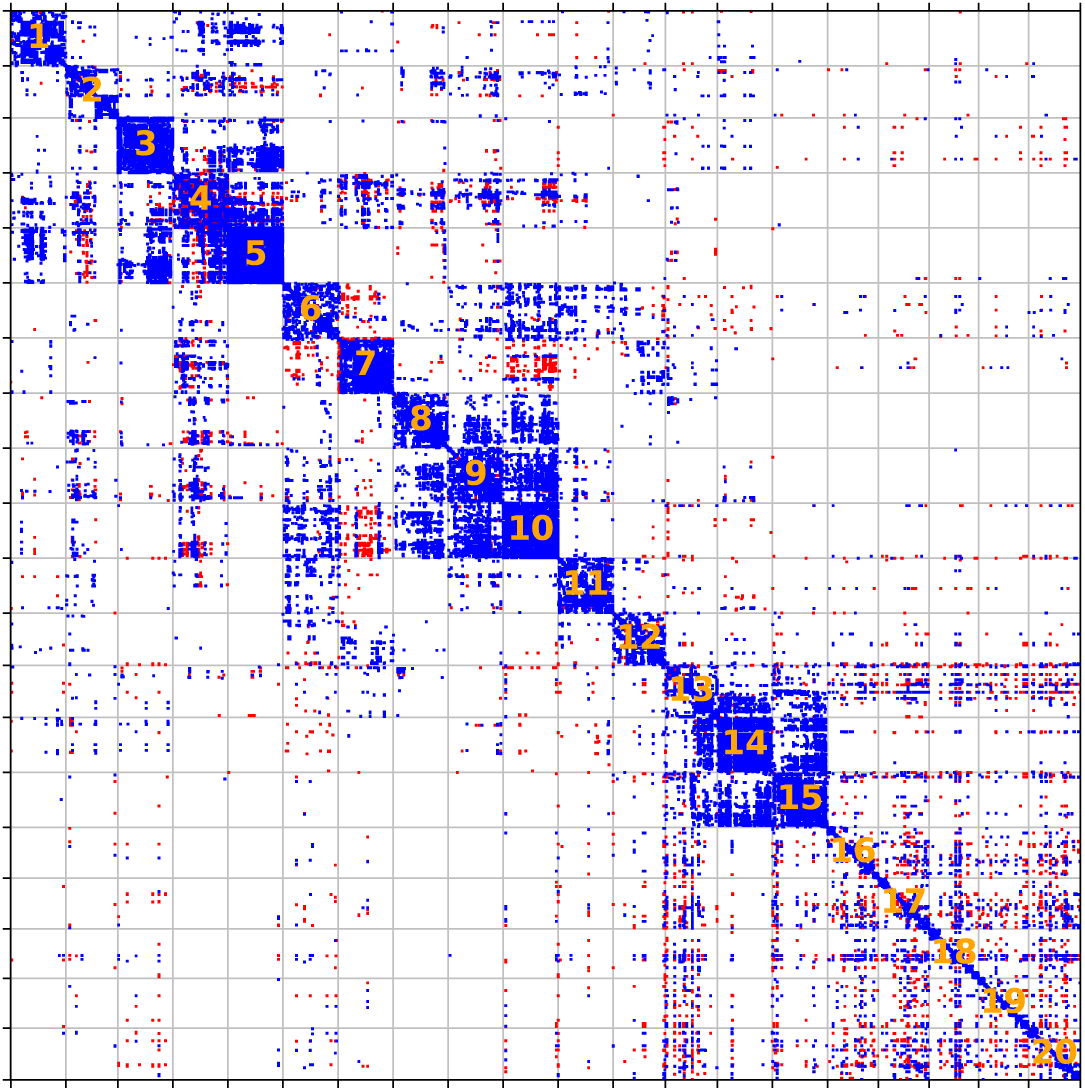
Asthma posterior gene network. The posterior gene network **Λ_y|x,Z_** after taking into account the phenotype data.

Thus, all of the modules that influence phenotypes, modules 13-20, showed enrichments in immune-related genes, with significant enrichment for modules 13-15 and weaker but still significant enrichment for modules 16-20. Since asthma is an immune disorder, the enrichment of immune-related genes in the trait-perturbing modules provides evidence that these modules are likely to play an important role in asthma patients.

#### SNPs perturbing asthma phenotypes overlap with SNPs perturbing immune modules

The SNP perturbation of the gene modules above (Figure 4C) may or may not result in a change in phenotypes. To see if the SNPs perturbing each gene module have an impact on the lung and blood phenotypes, we compared the top module-specific eQTLs in 0**Θ**_xy_ with the SNPs with the strongest effects on the lung or blood phenotypes in **B_xz_** inferred from our sparse Gaussian chain graph model. The SNPs with the strongest effects on the lung (or blood) phenotypes were determined based on the sum over the SNP effects on all lung (or blood) phenotypes in **|B_XZ_|.** Similarly, the top module-specific eQTLs were determined based on the sum over the SNP effect sizes on all expression levels in each module in |**Θ**_xy_| We obtained the overlap between the SNPs perturbing the phenotype network and the SNPs perturbing the gene network, considering the top 100 and 200 module-specific eQTLs and top 200 SNPs perturbing each phenotype group (the cutoff for top 200 SNPs shown as the magenta line at SNP effect size 0.013 for lung traits in Figure S1A and at SNP effect size 0.0037 for blood traits in Figure S1B). Using Fisher’s exact test, we also assessed the significance of these overlaps within the set of SNPs with non-zero effects in **Θ**_xy_.

In our comparison, only a subset of the eQTLs influenced phenotypes, but the eQTLs perturbing the immune modules, modules 13-20, were more likely to perturb the phenotypes than the eQTLs for the other modules (Figures 6A and 6B). Λmong the top 100 module-specific eQTLs, only a fraction of those SNPs overlapped with top 200 SNPs perturbing phenotypes (ranging from 0% to 18% of eQTLs across modules for an overlap with SNPs perturbing lung phenotypes and ranging from 2*%* to 20 % of eQTLs across modules for an overlap with SNPs perturbing blood phenotypes). These fractions increased as we considered more eQTLs as in top 200 and all module-specific eQTLs. This matches with the observations from previous studies that not all of the eQTLs affect higher-level phenotypes^21^ and that trait-associated SNPs are likely to be eQTLs ^40^.However, in our analysis, the eQTLs for immune-related modules, modules 13-20, tended to have larger overlaps than the other modules (Figures 6A and 6B). Furthermore, we found these overlaps are statistically significant for all of the immune modules, modules 13-20, but not for all of the other modules, and the most statistically significant overlaps were from the immune modules (Figures 6C and 6D). This suggests that eQTLs that perturb the modules that influence phenotypes are more likely to perturb phenotypes than eQTLs that perturb other gene modules.

**Figure 6.**
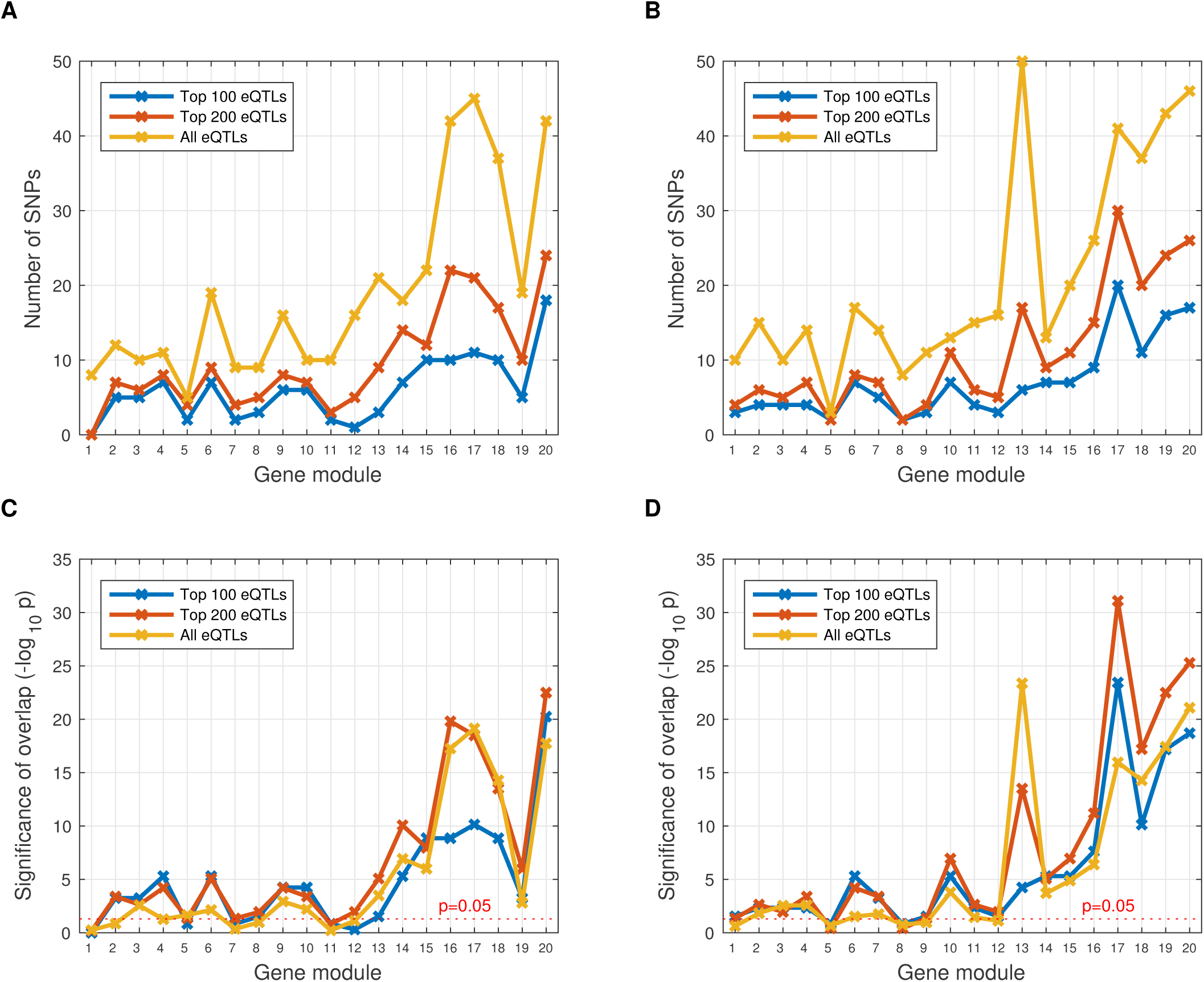
Overlap between SNPs perturbing phenotype network and SNPs perturbing gene network. For each gene module and each group of lung and blood phenotypes, we found the overlap between the top 200 SNPs perturbing the phenotype subnetwork and each of the top 100, 200, and all eQTLs perturbing the gene module. The number of SNPs in the overlap is shown for (A) lung phenotypes and (B) blood phenotypes. Statistical significance of the overlap is shown for (C) lung phenotypes and (D) blood phenotypes.

#### The immune modules mediate SNP perturbation of phenotypes

To understand the molecular mechanisms that underlie the SNPs perturbing phenotypes beyond the simple overlap of SNPs perturbing the phenotype network and SNPs perturbing the gene network, we used the Perturb-Net inference procedure to obtain the decomposition of the SNP effects on phenotypes **B**_xz_ into the component SNP effects on phenotypes 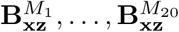 mediated by each of the 20 gene modules. We examined this decomposition for the 50 SNPs with the strongest effects on each group of lung and blood phenotypes (the cutoff for top 50 SNPs is shown as the green line at SNP effect size 0.04 for the lung phenotypes in Figure S1Λ and at SNP effect size 0.011 on blood phenotypes in Figure SIB),

For each set of 50 SNPs with the strongest perturbation effects on lung or blood phenotypes, nearly all of their effects on phenotypes were mediated by modules 12 through 20. The decomposition of the SNP effects on lung phenotypes (Figure 7A) into the 20 components (Figure 7B) shows that only the components for modules 12 through 20 contain non-zero SNP effects on the lung phenotypes, ex;cept for module 6, which mediates the effects of SNP rs1008932. We further summarized the component SNP effects by summing across all lung phenotypes for each SNP 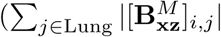 module *M* and SNP *i*; Figure 7C). In Figure 7C, for 45 out of the 50 SNPs the SNP effect on the lung phenotypes is mediated by a single module from modules 12-20. For the other 5 SNPs, although their effects on phenotypes were mediated by two or three modules, the module with the strongest mediator effect had effect size at least 5 times as large as the other modules. Although only 20 SNPs overlapped between the two sets of top 50 SNPs for lung and blood phenotypes, the SNP effects on blood phenotypes were also mediated by modules 12-20 (Figure.8), This indicates that modules 12-20 can potentially explain the molecular mechanisms behind the SNP perturbations of asthma phenotypes.

**Figure 7.**
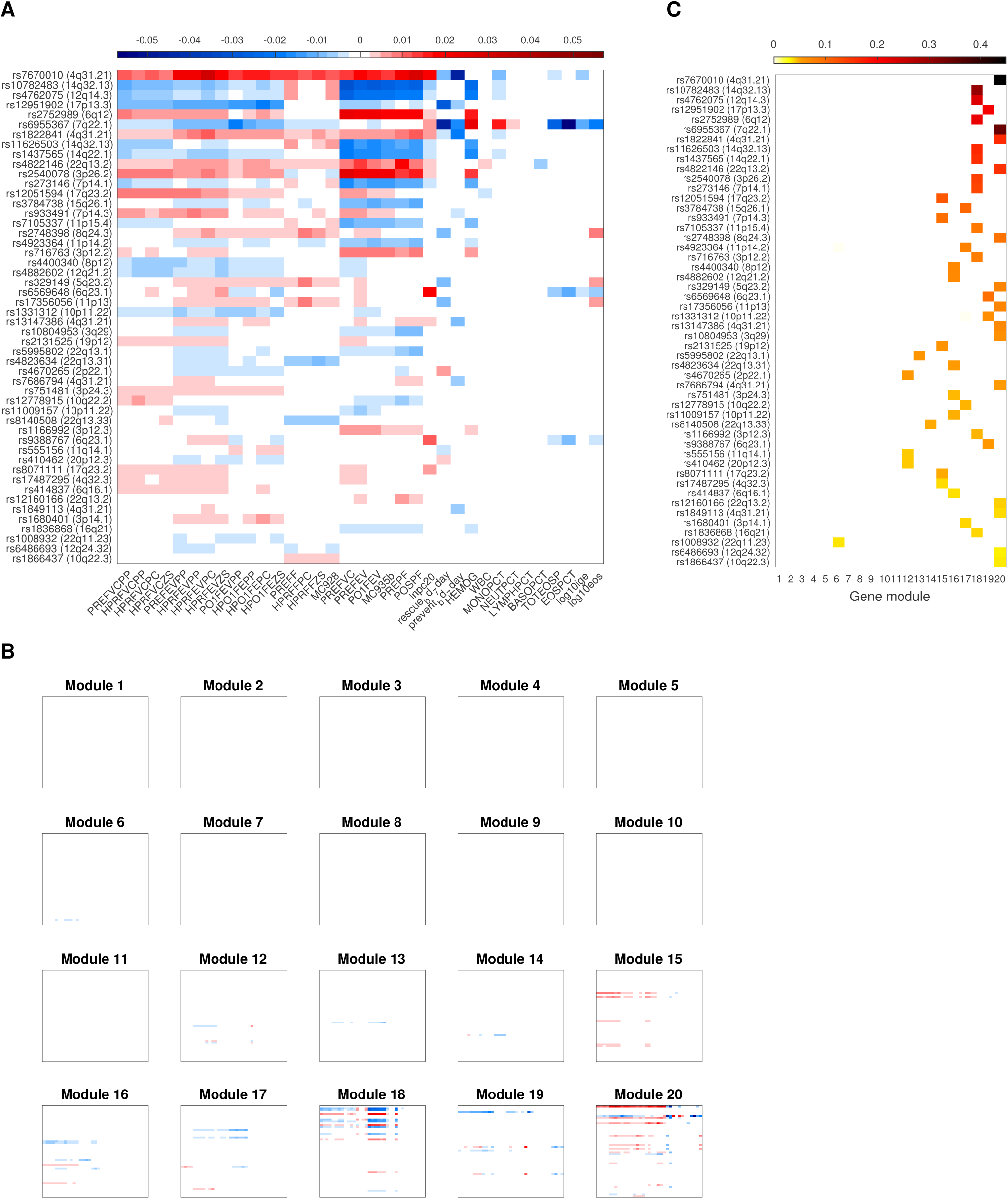
Top 50 SNPs perturbing lung phenotypes and their perturbations effects on phenotypes mediated by gene modules. For the top 50 SNPs perturbing lung phenotypes, we show (Λ) their effect sizes on phenotypes **B_xz_** and (B) the decomposition of **B_xz_** into component effects 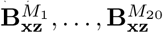 mediated by each of the 20 gene modules. The sum over all component effects in Panel (B) is equal to the overall effects in Panel (A). (C) We summarize each component SNP effect 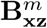 for module *m* in Panel (B) as a row-wise sum of 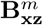, shown as the mth column in the figure. The SNPs are ordered according to their overall effect sizes on the lung phenotypes.

**Figure 8.**
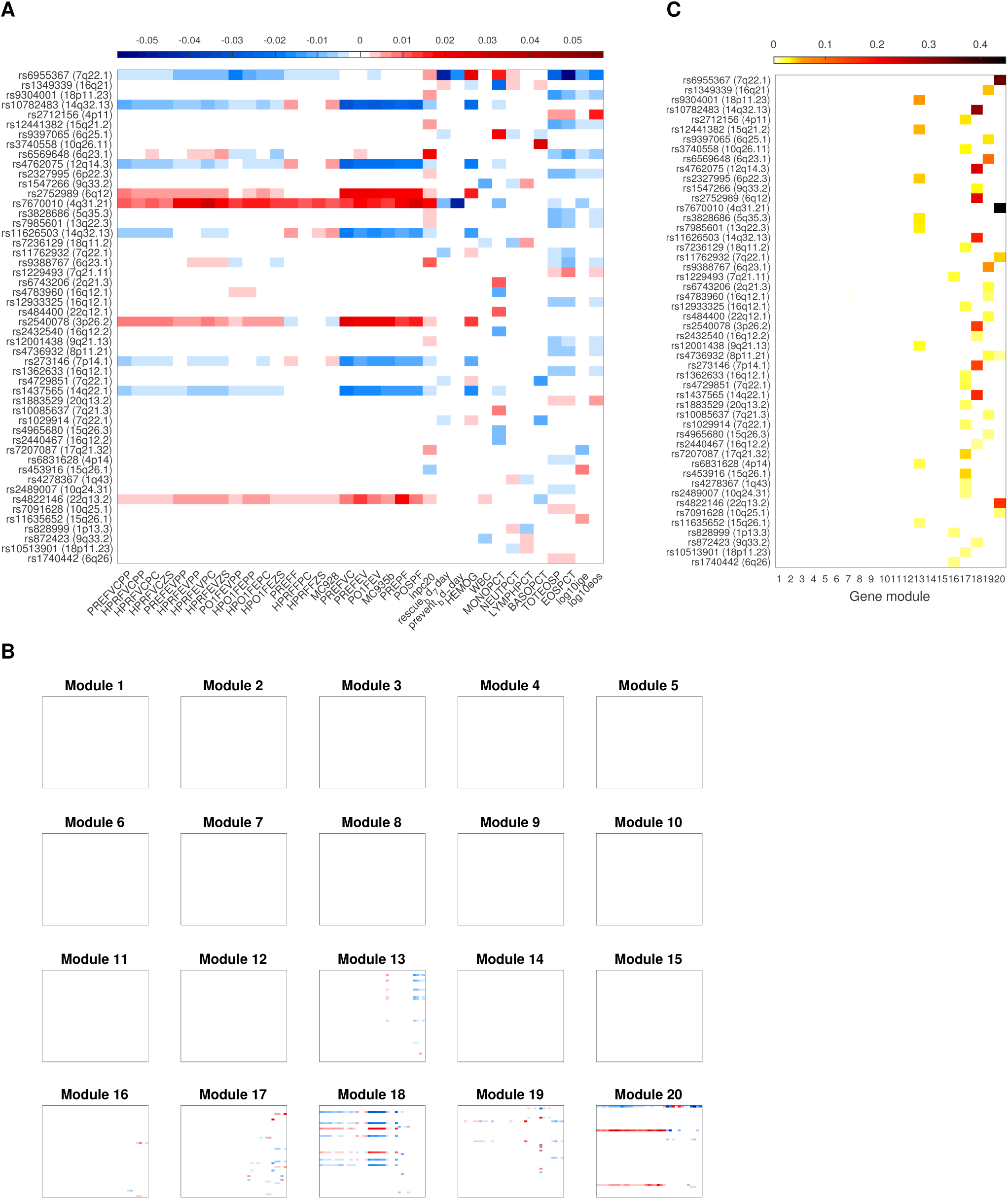
Top 50 SNPs perturbing blood phenotypes and their perturbations effects on phenotypes mediated by gene modules. For the top 50 SNPs perturbing blood phenotypes, we show (Λ) their effect sizes on phenotypes **B_xz_** and (B) the decomposition of **B_xz_** into component effects 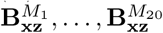 mediated by each of the 20 gene modules. The sum over all component effects in Panel (B) is equal to the overall effects in Panel (A). (C) We summarize each component SNP effect 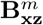 for module *m* in Panel (B) as a row-wise sum of 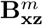, shown as the mth column in the figure. The SNPs are ordered according to their overall effect sizes on blood phenotypes.

#### Module 13 explains the molecular mechanism of the previously known association between SNP rs63340 and asthma susceptibility

We performed an in-depth analysis of module 13, its influence on asthma phenotypes, and its perturbation by SNP rs63340, one of the SNPs with the strongest effects on this module and also on phenotypes (Figure 9). SNP rs63340 ranked third for its effect on module 13 and 41st for its effect on phenotypes. The genome region 16q21, where SNP rs63340 is located, has been previously found to be linked to asthma and atopy in several previous genome-wide screenings,^41,42,43,44,45^ though the mechanism behind this association has not been fully elucidated. Our model indicated that this locus directly perturbs the expression levels of *NRP1, DCANP1, EPHB1, NLRP7* and *GZMB.* Several of these genes have been previously linked to asthma. *NRP1* is known to be a part of one of important mediators involved in the pathogenesis of asthma.^46^ Λ promoter nucleotide variant in *DCANP1* was previously associated with serum IgE levels among asthmatics. *EPHB1* has been previously linked to lung function traits in asthma.. ^48^ Our model found *EPHA2, KCNA5, NRP1*, and *CLEGC4C* as key mediator genes, determined by the row of 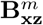 for SNP rs63340 and for gene *m* in module 13 summed across all phenotypes, that mediate the effects of SNP rs63340 on asthma phenotypes. Λmong these genes, *KGNA5* has been known to be connected to pulmonary vasoconstriction^49,50^ and a SNP near *KCNA5* was significantly associated with asthma.^51^ In addition, *CLEGC4C* has been known to be involved in immune response.^52,53^ Thus, the results from Perturb-Net are well supported by the previous findings in the literature and provide insights into the gene network underlying the previously reported association between the locus and asthma phenotypes.

**Figure 9.**
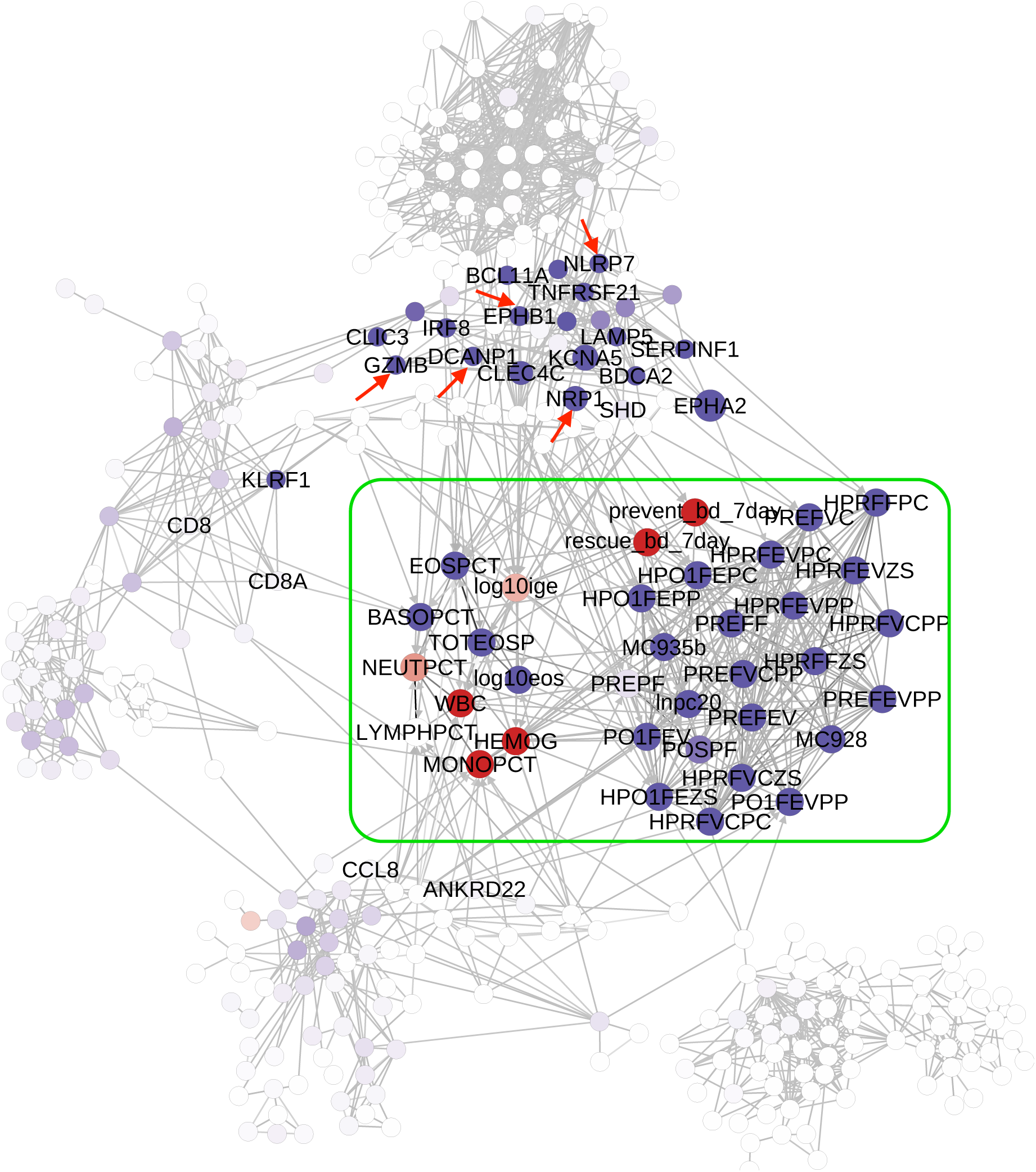
The gene network for module 13, its influence on asthma phenotypes, and its perturbation by SNP rs63340. The asthma phenotype network **Λ_z_** in our estimated model is shown in the green box and the gene network **Λ_y_** for module 13 is shown outside of the green box. The edges across the two networks correspond to direct influence of expression levels on phenotypes Θ_yz_. The five genes. (*NRP1, DCANP1, EPHB1, NLRP7, and. GZMB)* whose, expression is directly perturbed by SNP rs63340 with effect size > 0.05 in Θ**_xy_** are labeled with arrows, colored red to indicate positive eQTL effects. Node colors depict the indirect effects of this eQTL on gene expression levels **B_xy_** and phenotypes **B_xz_,** with red for up-regulation and blue for down-regulation. Node size: of genes depicts the component of the eQTL effects on phenotypes mediated by the. given gene, the row of 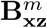 for SNP rs63340 and for gene *m* in module 13 summed across all phenotypes.

### Comparison with other methods

We compared our method with the two-layer Lasso both qualitatively by visual inspection of the estimated parameters and quantitatively by assessing the predictive power of different methods.

#### Comparison of the estimated models

We compared the results from our approach and the two-layer Lasso by visually inspecting the estimated SNP effects on gene modules and the estimated gene module effects on phenotypes. For the top 50 SNPs perturbing the lung phenotypes (Figure 7), we examined the overall SNP effects on each gene module based on Θ_xy_ and B_xy_ from our model and **A**_xy_ from the two-layer Lasso (Σ_*j*∈M_|[**Θ**_xy_]_*i,j*_|,Σ_*j*∈M_|[**B**_xy_]_*i,j*_|,and Σ_*j*∈M_|[**A**_yz_]_*j*,k_| for SNP *i* module *M*) To see how each gene module influences phenotypes, we computed the magnitudes of overall gene module effects on each phenotype from **Θ**_yz_ and **B**_yz_ in our model and **A**_yz_ in the two-layer Lasso (Σ_*j*∈M_|[**Θ**_yZ_]_*j*,k_|,Σ_*j*∈M_|[**B**_yz_]_*j*,k_|,and Σ_*j*∈M_|[**A**_yz_]_*j*,k_| for module *M* and phenotype *k*).

Unlike the Perturb-Net model, the two-layer Lasso does not model direct and indirect perturbation effects separately but attempts to capture both types of effects in a single set of parameters. Thus, the perturbation effects captured by the two-layer Lasso appeared to be a compromise between the direct and indirect perturbation effects captured by Perturb-Net (Figure 10). However, the SNP effects appeared to be similar across **Θ**_xy_, **B**_xy_, and **A**_xy_ in the module-level summaries (Figures l0A-l0C), because the direct SNP perturbation effects tended to propagate to other genes only within each module, but not to genes in other modules. On the other hand, the module effects on phenotypes showed a distinct pattern across **Θ**_yz_, B_yz_, and A_yz_ (Figures 10D-10F), because in our model, the direct influence of gene expression levels on a phenotype induces the indirect influence on other correlated phenotypes, whereas the Lasso parameter tries to capture both types of information in a single parameter.

**Figure 10.**
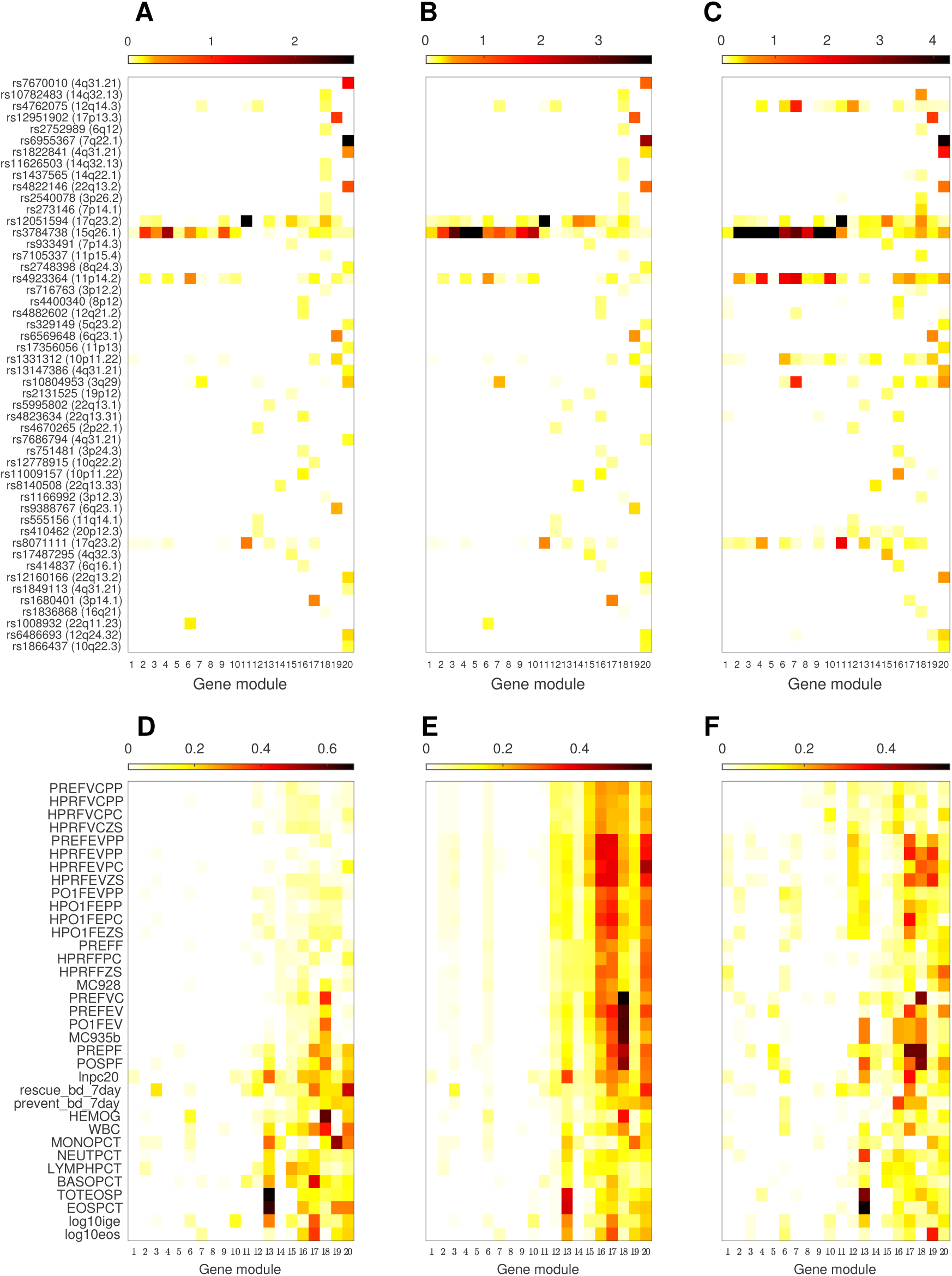
Comparison of different methods for learning the cascaded influence of SNPs to gene modules to phenotypes. For the top 50 SNPs perturbing lung phenotypes (Figure 7), the effects of these SNPs on each of the gene modules are shown for (A)Θ_yz_ from our model, (B) **B_yz_** inferred from our model, and (C) **Λ_yz_** from the two-layer Lasso. The effects of the expression levels in each gene module on lung phenotypes are shown for (D)Θ_yz_ from our model, (E) **B_yz_** inferred from our model, and (F)**A_yz_** from the two-layer Lasso. The effect sizes in each model parameter matrix above were summed across all genes within each module after taking absolute values.

#### Comparison of prediction accuracy

We assess the ability to make predictions about new asthma patients based on the estimated Perturb-Net model and compare the results with those from the two-layer Lasso. We split the data into train and test sets and obtained the prediction accuracy using the test set after training a model on the train set. We set aside 25 samples as a test set and used the remaining 115 fully-observed samples and 34 partially observed samples to train a sparse Gaussian chain graph model with our semi-supervised learning method. We also trained a model, using only the 115 fully observed samples with the supervised learning method, and compared the results from the two-layer Lasso, also trained from the fully observed samples. Given the estimated models, we performed prediction tasks and obtained the prediction error as the squared difference between the observed and predicted values averaged across samples in test set.

The Perturb-Net model estimated from all data had the smallest prediction error for all of the prediction tasks (Table 2). In particular, our model with semi-supervised learning performed better than our model with supervised learning, demonstrating that leveraging partially observed data can help learn a model with greater predictive power. For supervised learning, our model outperformed Lasso. This demonstrates that taking into account the network structure in expression levels and clinical phenotypes increases the performance on prediction tasks.

**Table 2.**
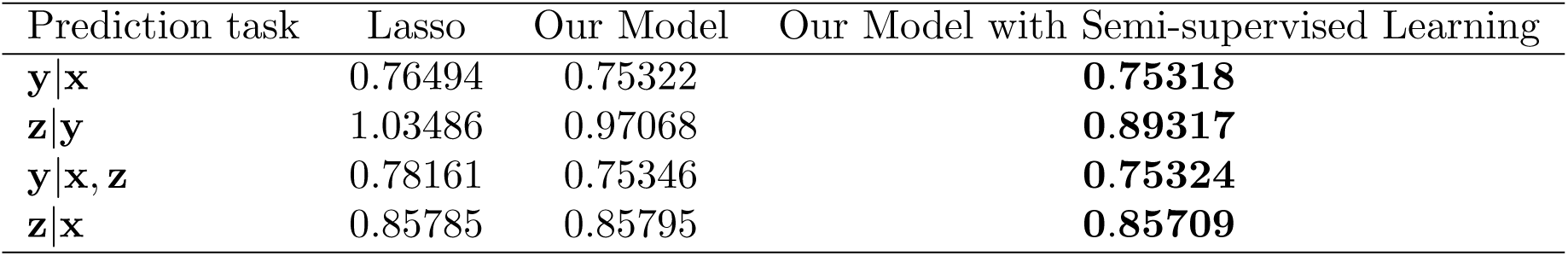
Prediction errors of different methods on asthma test set

## Discussion

We introduced a statistical framework called Perturb-Net for learning a gene network underlying phenotypes using SNP perturbations and for identifying SNPs that perturb this network, given population genotype, expression, and phenotype data* Compared to many of the previous methods that focused on the co-localization of eQTLs and genetic association signals for phenotypes,^13,14,15^ using multi-stage methods,^10,11,12^ our approach combines all available data in a single statistical analysis and directly models the multiple layers of a biological system with a. cascade of influence from SNPs to expression levels to phenotypes, while modeling each layer as a network. Our probabilistic graphical model framework allows to model eQTLs with or without an impact on phenotypes for an investigation of co-localization of SNPs perturbing expression levels and SNPs perturbing phenotypes and to extract rich information on the molecular mechanisms that explains the influence of SNPs on phenotypes. We developed fast learning algorithms called Fast-sCGGM and Mega-sCGGM for learning sCGGM components of the Perturb-Net model, which serve as the key subroutine of our Perturb-Net learning method, to enable analysis of human genome scale data within a few hours.

Our results from applying Perturb-Net to asthma data, confirmed the observations from the previous studies, including GWΛS, eQTL mapping, and gene network modeling,^11,40^ Our results confirmed the finding from previous studies on combining the results of GWΛS and eQTL mapping^40^ that there is a. partial overlap between SNPs perturbing expression levels and SNPs perturbing phenotypes. In addition, this overlap was more significant for eQTLs that perturb trait-associated modules than eQTLs that perturb other parts of the gene network, as was previously reported.^21^

The analysis of the asthma data with Perturb-Net provided new insights. Perturb-Net was able to systematically reveal the gene network that lies between the SNPs and phenotypes and to uncover how different parts of this gene network modulate the SNP effects on phenotypes in a statistically principled manner. Often, there are genetic loci that have been previously known to be linked to the disease susceptibility, though little is known about the underlying molecular mechanism. In such cases, the Perturb-Net analysis of asthma data demonstrated the potential to reveal the molecular pathway that are perturbed by previously known trait-associated loci.

Perturb-Net provides a flexible tool that can be extended in several different ways in a straightforward manner. Because the sparse Gaussian chain graph model in Perturb-Net uses sCGGMs as building blocks, the sCGGM component models can be threaded in different ways to form sparse Gaussian chain graph models with different structures. For example, if expression data from multiple tissue types are available for a patient cohort along with genome sequence and phenotype data, a sparse Gaussian chain graph model can be set up with multiple component sGGGMs_t_ each corresponding to the gene network under SNP perturbation in each tissue type, linked to another sCGGM for modeling expression levels influencing phenotypes. Models like this can reveal SNPs that perturb phenotypes through different tissue types and through different modules in each tissue-specific gene network. Λnother possible extension is to thread more than two component sCGGMs within a sparse Gaussian chain graph model to model more than two layers in a biological system, including epigenomes, metabolomes, and proteomes.

## Appendix A: Fast-sCGGM and Mega-sCGGM for efficiently learning sCGGMs

We introduce our scalable learning algorithms for sCGGM, since the learning algorithm for sparse Gaussian chain graph models in Eq. (1) uses the sCGGM learning algorithm as a key module. Assume an sCGGM^7,8^ for gene expression levels ye∈ℝ^*q*^ for *q* genes and genotype data x ∈ *£* {0,1,2}^*p*^ for minor allele frequencies at *p* loci be given as follows:

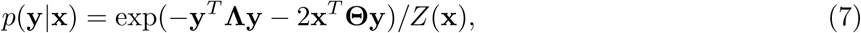

where **Λ** is a *q× q* matrix representing a. gene network, Θ is a *p ×q* matrix modeling SNPs influencing the expression levels of genes in the network, and *Z*(x) *=* (2π)^*q*/2^|Λ|^_1^ exp(x^*T*^**Θ0Λ**^-1^**Θ** ^T^x) is the constant to ensure the probability distribution integrates to 1. Then, given genotype data X ∈ℝ^*n*×*p*^ for *n* samples and *p* SNPs, each element taking a value from {0,1, 2} for the number of minor alleles at the locus, and expression data. Y∈ℝ^*n*×*q*^ for *q* genes for the same samples, a parameter estimate of the sCGGM in Eq. (7) can be obtained by minimizing Li-regula.rized negative log-likelihood:

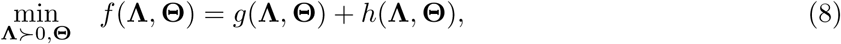

Where *g*(**Λ**,**Θ**)=-log |Λ|+(**S**_yy_**Λ**+2S_xy_^*T*^**Θ** 0+Λ^-1^_^1^Θ^*T*^S_xx_**Θ**) is the smooth negative log-likelihood, given data covariance matrices 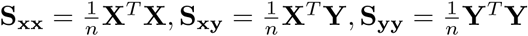, and *h*(Λ,Θ)= λ_Λ_|| Λ ||_1_+λ _Θ_||Θ||_1_ for the non-smooth elementwise *L*_1_ penalty. λ _Λ_a, λ_Θ_> 0 are regularization parameters.

Below, we introduce Fast-sCGGM for learning an sCGGM that substantially reduces computation time by orders of magnitude compared to the previous sta.te-of-the-a.rt method.^22^ Then, we describe Mega-sCGGM, a modification of Fast-sCGGM, that performs block-wise computation to learn a model from large human genome-wide data on a. machine with limited memory.

### Fast-sCGGM for improving computation time

Fast-sCGGM uses an alternate Newton coordinate descent method that alternately updates **Λ** and 0Θ, optimizing Eq. (8) over **Λ** given **Θ** and vice versa, until convergence. Our approach is based on the key observation that with Λ fixed, the problem of solving Eq. (8) over **Θ** becomes simply the well-known Lasso optimization, which can be solved efficiently using a. coordinate descent method.^54^

On the Other hand, optimizing Eq. (8) for **Λ** given **Θ** requires forming a quadratic approximation to find a generalized Newton direction and performing line search to find the step size. However, this computation is significantly simpler than performing the same type of computation on both **Λ** and **Θ** jointly as in the previous approach.^22^ Our algorithm iterates between the following two steps until convergence:

- **Coordinate descent optimization for Θ given Λ**: With Λ fixed, the optimization problem in Eq. (8) becomes

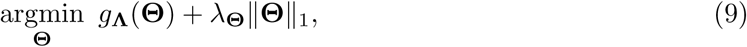

where *g*_Λ_(**Θ**)=tr(2S_xy_^*T*^**Θ**+Λ^-1^**Θ**^*T*^S_xx_**Θ**). Since *g*_Λ_(**Θ**) is a quadratic function, Eq. (9) corresponds to the Lasso problem and the coordinate descent method can be used to solve this efficiently.
- **Coordinate descent optimization for** Λ **given Θ**: Given fixed **Θ**, the problem in Eq, (8) becomes

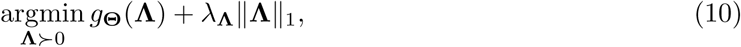

where *g***_Θ_**(Λ)=- log |Λ| +tr(S_yy_Λ+Λ^-1^**Θ**^*T*^S_xx_**Θ**). In order to solve this, we first find a generalized Newton direction that; minimizes the L_1_-regularized quadratic approximation 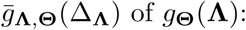:

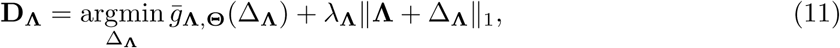

where 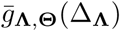 is obtained from a second-order Taylor expansion and is given as

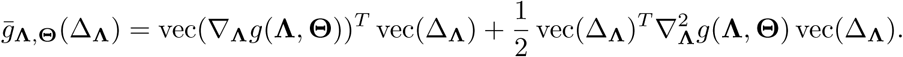 In the above equation,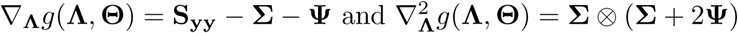, where **Σ**=Λ^-1^ and Ψ=**ΣΘ**^*T*^S_xx_**ΘΣ**, are the components of the gradient and Hessian matrices corresponding to **Λ**. The problem in Eq. (11) is again equivalent to the Lasso problem, which can be solved efficiently via coordinate descent. Given the Newton direction for **Λ**, we update **Λ←Λ**+*α*D**_Λ_**, where step size 0 <*α≤*1 ensures sufficient decrease in Eq. (8) and positive definiteness of **Λ**. The *α* is obtained by line search on the objective in Eq. (10). In order to further reduce computation time, we adopt the following strategies that have been previously used for sparse Gaussian graphical model and sCGGM optimizations.^22,55^ First, to improve the efficiency of coordinate descent for the Lasso problem in Eqs. (9) and (11), we restrict the updates to an active set of variables given as:

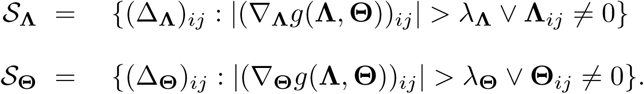 Because the active set sizes *m***_Λ_**=|*S***_Λ_**|,*m***_Θ_**= |*S* Θ| approach the number of non-zero entries in the sparse solutions for **Λ** and **Θ** over iterations, this strategy yields a substantial speedup. Second, to further improve the efficiency of coordinate descent, we store intermediate results for the large matrix products that need to be computed repeatedly. We compute and store **U**:=Δ_Λ_Σ and V:=Δ_Θ_Σ at the beginning of the optimization. Then, after a coordinate descent update to (Δ_Λ_)_*i,j*_, row *i* and *j* of **U** are updated. Similarly, after an update to (Δ_Θ_)_*i,j*_, row *i* of **V** is updated. Finally, in each iteration of Fast-sCGGM, we warm-start **Λ** and **Θ** from the results of the previous iteration and make a single pass over the active set. This ensures decrease in the objective in Eq. (8), while reducing the overall computation time in practice. The pseudocode for Fast-sCGGM is provided in Λlgorithm 1.

### Mega-sCGGM for removing memory requirement

Fast-sCGGM as described above is still limited by the space required to store large matrices during coordinate descent computation. Solving Eq. (11) for updating **Λ** requires precomputing and storing *q×q* matrices,Σand Ψ=**ΣΘ**^*T*^S_xx_**ΘΣ**, whereas solving Eq.(9) for updating **Θ** requires **Σ** and a *p×p* matrix S_xx_. A naive approach to reduce the memory footprint would be to recompute portions of these matrices on demand for each coordinate update, which would be very expensive.

#### Algorithm 1 Fast-sCGGM

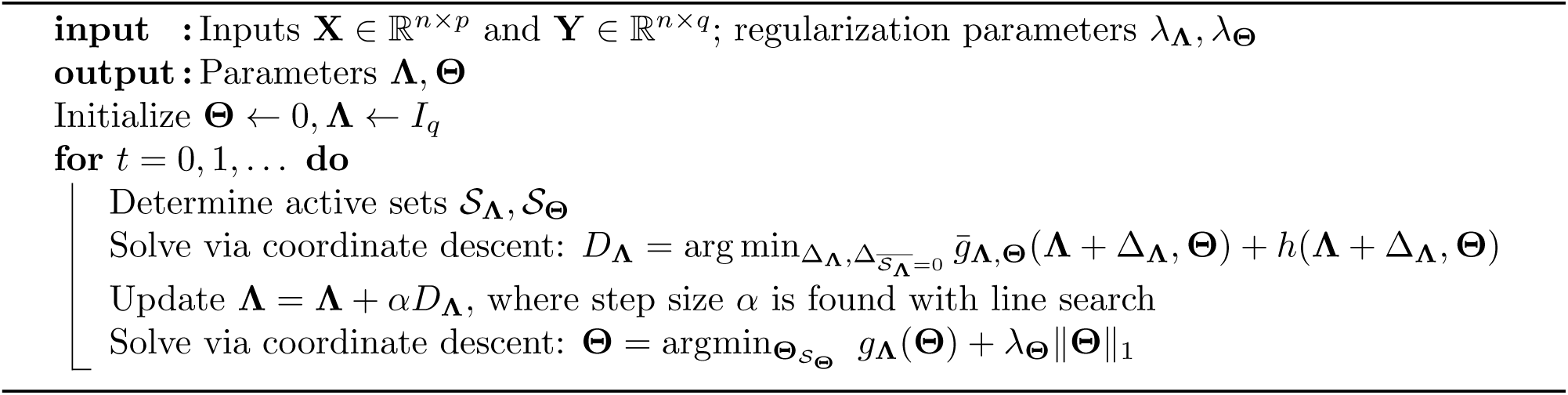

Here, we describe Mega-sCGGM that combines the alternating Newton coordinate descent algorithm in Fast-sCGGM with block coordinate descent to scale up the optimization to very large problems on a machine with limited memory. During coordinate descent optimization, we update blocks of **Λ** and **Θ** so that within each block, the computation of the large matrices can be cached and re-used, where these blocks are determined automatically by exploiting the sparse stucture. For **Λ**, we extend the block coordinate descent approach in BigQUIC^56^ developed for sparse Gaussian graphical models to take into account the conditioning variables in CGGMs. For **Θ**, we describe a new approach for block coordinate descent update. Our algorithm can, in principle, be applied to problems of any size on a machine with limited memory and converges to the same optimal solution as our alternating Newton coordinate descent method.

#### Blockwise Optimization for Λ

A coordinate descent update of [Δ_Λ_]_*i,j*_, requires the *i*th and *j*th columns of **Σ** and Ψ If these columns are in memory, they can be re-used. Otherwise, it is a cache miss and we should compute them on demand as follows. We obtain [**Σ**]:, _*i*_ by solving linear system Λ[**Σ**]:,_*i*_= e_*i*_, where e_*i*_ is a vector of *q* 0’s except for 1 in the ith element, with conjugate gradient method. Then,[Ψ]:,_*i*_can be obtained from R^*T*^ [R]:,_*i*_, where R = X**ΘΣ**.

In order to reduce cache misses, we perform block coordinate descent, where within each block, the columns of **Σ** are cached and re-used. Suppose we partition *N* = {1, …, *q*} into *k***_Λ_** blocks,

##### Algorithm 2 Mega-sCGGM

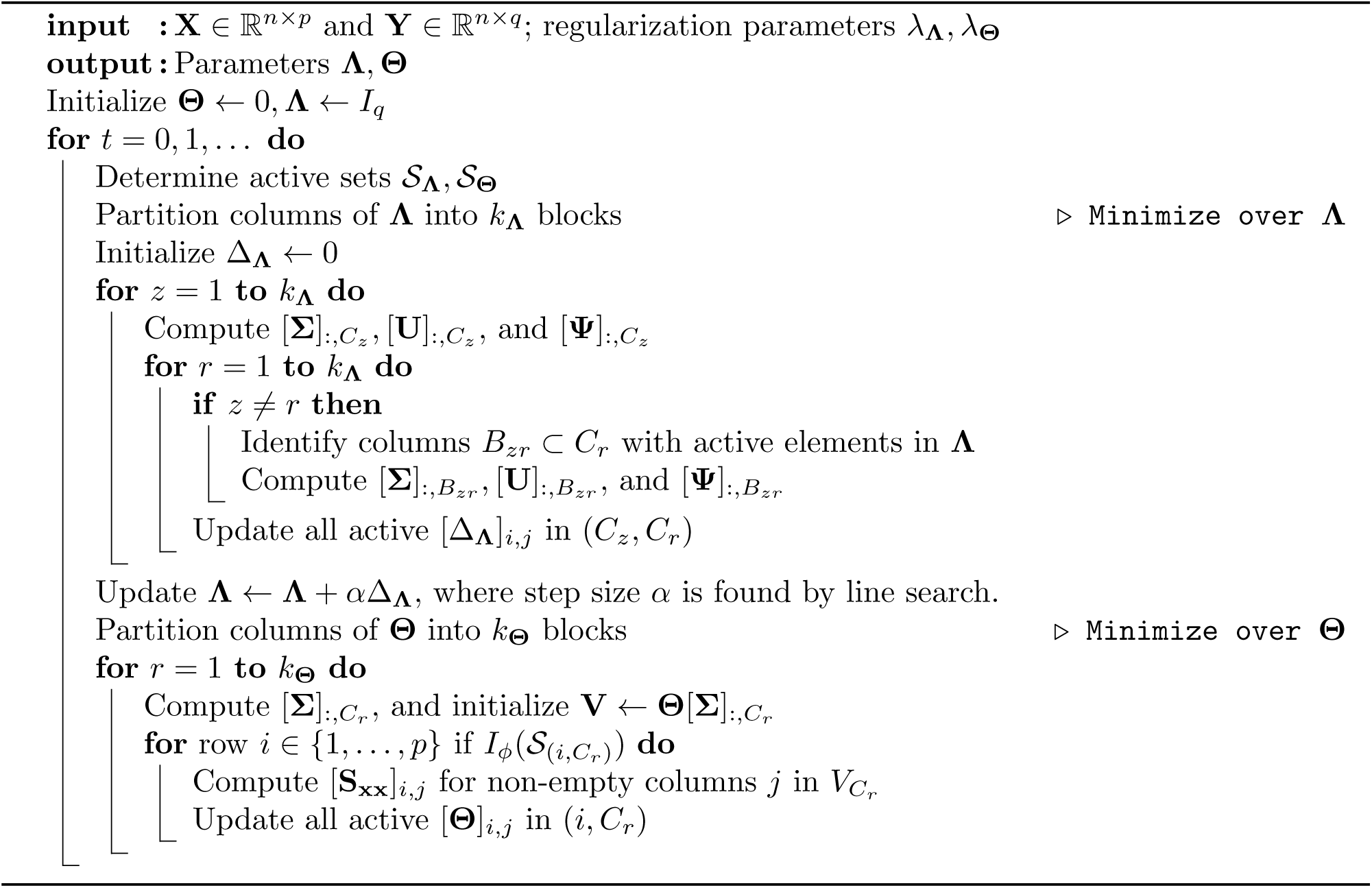

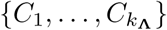. We apply this partitioning to the rows and columns of Δ**_Λ_** to obtain *K***_Λ_** x *K***_Λ_** blocks. We perform coordinate-descent updates in each block, updating all elements in the active set within that block. Let 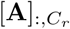 denote a matrix containing columns of **A** that correspond to the subset *C*_*r*_. In order to perform coordinate-descent updates on *(C*_*r*_, *C*_*z*_) block of Δ **_Λ_,** we need 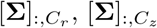, 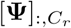, and 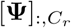 Thus, we pick the smallest possible *K***_Λ_** such that we can store *2q/ K***_Λ_** columns of **Σ** and **Ψ** in memory. When updating the variables within block *(C*_*z*_, *C*_*r*_) of Δ**_Λ_**, there are no cache misses once 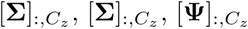, and 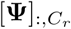 are computed and stored. **Λ**fter updating each [Δ**_Λ_**]_*i,j*_ to [Δ**_Λ_**] _*i,j*_ + *μ,* we maintain 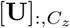 and 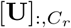 by

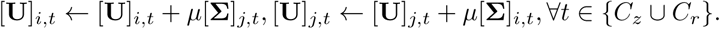

To go through all blocks, we update blocks *(C*_*z*_, *C*_1_*),(C*_*z*_, *C*_*k*_) for each *z ϵ* {1,…, *K***_Λ_** }. Since all of these blocks share 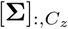 and 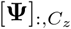, we precompute and store them in memory. When updating an off-diagonal block (*C*_*z*_,*C*_*r*_*),z* ≠ r, we compute 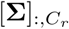 and 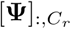 Overall, each block of **Σ** and **Ψ** will be computed *K***_Λ_** times.

In typical real-world problems, the graph structure of **Λ** will exhibit clustering, with an approximately block diagonal structure. We exploit this structure by choosing a partition 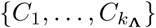 that reduces cache misses. Within diagonal blocks (*C*_*r*_, *C*_*r*_)*’*s, once 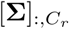 and 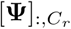 are computed, there are no cache misses. For off-diagonal blocks (*C*_*r*_, *C*_*z*_)*’*s,*r ≠ z,* we have a cache miss only if some variable in {[Δ]_*i,j*_ | *i ϵ C*_*r*_, *j ϵ C*_z_} lies in the active set. We minimize the active set in off-diagonal blocks via clustering, following the strategy for sparse Gaussian graphical model estimation in BigQUIC^56^ and using the METIS^37^ graph clustering library.

Although the worst-case scenario is to compute **Σ** and **Ψ** *K***_Λ_** a times to update all elements of Δ**_Λ_**, in practice, graph clustering dramatically reduces this additional cost of block-wise optimization. In the best case, if the active set for **Λ** is perfectly block-diagonal and graph clustering identifies this block diagonal structure, we need to compute **Σ** and **Ψ** only once to update all the blocks. A depiction of our blockwise optimization scheme is given in Figure S2.

### Blockwise Optimization for Θ

The coordinate descent update of [Θ]_*i,j*_ requires [**S_xx_**]_: *i*_ and [**Σ**]_*j*_ to compute 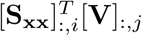 where [V]:,_*j*_ = **Θ**[**Σ**] _*j*_. If [**S_xx_**]_: *i*_ and [**Σ**] _*j*_ are not already in the memory, it is a cache miss. Computing [**S_xx_**]_:***i***_ takes *O*(*np*), which is expensive if we have many cache misses.

We propose a block coordinate descent approach for solving Eq. (9) that groups these computations to reduce cache misses. Given a partition {1,…, *q*} into *K*_Θ_ subsets, 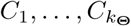, we divide **Θ** into *p* x *K*_Θ_ blocks, where each block comprises a portion of a row of **Θ**. We denote each block (*i,C*_*r*_), where *i ϵ* {1,…,*p*}. Since updating block (*i,C*_*r*_) requires [**S_xx_**]**:,_*i*_**.and 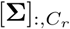, we pick smallest possible *K*_Θ_ such that we can store *q/ K*_Θ_ columns of **Σ** in memory. While performing coordinate descent updates within block (_*i*_ *C*_*r*_) of **Θ**, there are no cache misses, once [**S_xx_**]**_: *i*_** and 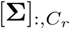, are in memory After updating each [**Θ**]_*i.j*_ to [**Θ**]_*i.j*_ + *μ* we update 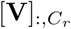 by 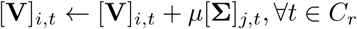

In order to sweep through all blocks, each time we select a *q ϵ* {1,…, *K*_Θ_ } and update blocks (1, *C*_*r*_),…, (*p, C*_*r*_). Since all of these *p* blocks with the same *C*_*r*_ share the computation of 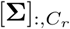, we compute and store 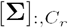 memory. Within each block, the computation of [S_xx_]_;,*i*_ is shared, so we precompute and store it in memory, before updating this block. The full matrix of Σ will be computed once while sweeping through the full **Θ**, whereas S_xx_ will be computed *K***_Θ_** times.

We further reduce cache misses for [S_xx_]_;,*i*_ by strategically selecting partition 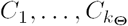, based on the observation that if the active set is empty in block (*i, C*_*r*_), we can skip this block and forgo computing [S_xx_]_;,*i*:_ We therefore choose a partition where the active set variables are clustered into as few blocks as possible. Formally, we want to minimize 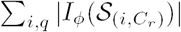 where 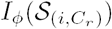 is an indicator function that outputs 1 if the active set 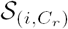 within block (*i,C*_*r*_) is not empty and 0 otherwise. We therefore perform graph clustering over the graph *G =* (V, *E)* defined from the active set in **Θ**, where *V =* {*1,…,q*} with one node for each column of **Θ**, and *E =* {(*j. k*)| [**Θ**]_*i,j*_, ϵ *𝒮* **_Θ_**, [**Θ**]_*i,k*_ ϵ *𝒮* _**Θ**_ for *i=* 1,….,*p*}, connecting two nodes *j* and *k* with an edge if both [**Θ**]_*i,j*_ and [**Θ**]_*i,k*_ are in the active set. This edge set corresponds to the non-zero elements of **Θ ^*T*^ Θ,** so the graph can be computed quickly in *O*(*m* **Θ** *q*).

We also exploit row-wise sparsity in **Θ** to reduce the cost of each cache miss. Every empty row in **Θ** corresponds to an empty row in V = **ΘΣ**. Because we only need elements in [S_xx_]_:_,_*i*_ for the dot product 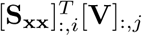, we skip computing the fcth element of [S_xx_]:,_*i*_ if the *k*th row of **Θ** is all zeros. Our blockwise optimization scheme for **Θ** is depicted in Figure S3.

### Parallelization in Fast-sCGGM and Mega-sCGGM

We parallelize some of the expensive computations in Fast-sCGGM and Mega-sCGGM on multi-core machines. For both methods, we parallelize all matrix-matrix and matrix-vector multiplications. In addition, we parallelize the computation of columns of **Σ** and **Ψ** in Fast-sCGGM and the computation of multiple columns of **Σ** and **Ψ** within each block in Mega-sCGGM. In Mega-sCGGM, we parallelize the computation of each row of S_xx_ whenever it is recomputed.

## Appendix B: Efficient implementation of EM algorithm for semi-supervised learning for Perturb-Net model

The standard EM algorithm iterates between an M-step for finding the parameter estimate maximizing the expected log-likelihood (or minimizing the negative log-likelihood) and an E-step for finding the expected sufficient statistics based on the posterior probability distribution in Eq. (5). The M-step is carried out by using our Mega-eCGGM algorithm. In E-step, a naive inversion of 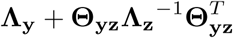 to obtain Σ_y|x z_ is expensive and storage of this dense matrix may exceed computer memory for large gene expression datasets. We reduce the time cost and avoid memory limit in E-step, assuming that the number of phenotypes *r* is relatively small compared to the number of genes (i.e., *r<<q*), which is typical for most studies. Instead of explicitly performing the E-step, we embed the E-step within the M-step, such that the E-step results are represented implicitly to fit in memory and computed explicitly on-demand as needed in the M-step. Specifically, instead of performing the full E-step, we implicitly represent 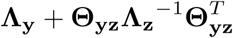 as **Λ**_y_ +**KK**^*T*^,using low-rank component 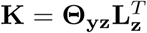 and the sparse Cholesky factorization of trait network 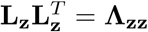. Then, during M-step, we invert Λ_y_ + **KK**^*T*^, one column at a time as needed, using the conjugate gradient method. This modified EM algorithm is equivalent to the original EM algorithm that iterates between an M-step and an E-step, producing the same estimate.

## Supplemental Data

Supplemental Data include 3 figures and 1 table.

## Acknowledgements

SYK, CM were supported by NSF CΛREER Λward No. MCB-1149885 (nsf.gov) and PΛ CURE (http://www.health.pa.gov/). The funders had no role in study design, data collection and analysis, decision to publish, or preparation of the manuscript.

## Declaration of Interests

The authors declare no competing interests.

## Web Resources

Perturb-Net software will be available at https://github.com/sssykim/Perturb-Net.

